# Repressor element 1-silencing transcription factor deficiency yields profound hearing loss through K_v_7.4 channel upsurge in auditory neurons and hair cells

**DOI:** 10.1101/2022.03.03.482811

**Authors:** Haiwei Zhang, Hongchen Li, Mingshun Lu, Shengnan Wang, Xueya Ma, Fei Wang, Jiaxi Liu, Xinyu Li, Haichao Yang, Fan Zhang, Haitao Shen, Noel J. Buckley, Nikita Gamper, Ebenezer N. Yamoah, Ping Lv

## Abstract

Repressor element 1-silencing transcription factor (REST) is a transcriptional repressor that recognizes neuron-restrictive silencer elements in the mammalian genomes in a tissue- and cell-specific manner. The identity of REST target genes and molecular details of how REST regulates them are emerging. We performed conditional null deletion of *Rest* (cKO) in murine hair cells (HCs) and auditory neurons, spiral ganglion neurons (SGNs). Null-deletion of full-length REST resulted in normal HCs and SGNs development but manifested with progressive hearing loss in adulthood. We found that deletion of REST results in an increased abundance of K_v_7.4 channels at the transcript, protein, and function levels. Specifically, we found that SGNs and HCs from *Rest* cKO mice displayed increased K_v_7.4 expression and augmented K_v_7 currents; SGN’s excitability was also significantly reduced. Administration of K_v_7.4 channel activator, fasudil, recapitulated progressive hearing loss in mice.

In contrast, inhibition of the K_v_7.4 channel by XE991 rescued the auditory phenotype of *Rest* cKO mice. Previous studies identified some loss-of-function mutations within the K_v_7.4-coding gene, *Kcnq4*, as a causative factor for progressive hearing loss in mice and humans. Thus, the present findings revealed that a critical homeostatic K_v_7.4 channel level is required for proper auditory functions.

## Introduction

The repressor element-1 silencing transcription factor (REST), also known as neuronal restriction silencing factor (NRSF), is a transcriptional repressor that binds a specific 21bp consensus sequence named repressor element 1 (RE-1) (Chong, Tapia-Ramirez et al., 1995, Schoenherr & Anderson, 1995). Through its DNA-binding domain, REST recruits multiple chromatin remodeling factors to form a repressor complex, ultimately repressing the transcription of target genes(Ooi & Wood, 2007, Schoenherr & Anderson, 1995). REST is initially expressed in embryonic stem cells, neural stem cells, and non-neural cells during embryogenesis. It maintains embryonic stem cells’ pluripotency and promotes stem cell differentiation and neuronal development, which are essential for neuronal diversity, plasticity, and survival (Chen, Paquette et al., 1998, Sun, Greenway et al., 2005). In non-neuronal cells, REST maintains a non-neuronal cell gene expression pattern by suppressing the expression of neural-related genes through histone deacetylation, chromatin remodeling, and methylation^(Chen et al., 1998)^. Recent studies suggest that the role of REST expands beyond neuronal development. Nearly 2000 genes contain predicted REST binding sites (Bruce, Donaldson et al., 2004, Johnson, Mortazavi et al., 2007, Seki, Masaki et al., 2014), and up to 90% of these REST target genes are tissue-and cell-type-specific (Bruce, Lopez-Contreras et al., 2009, Hohl & Thiel, 2005). As a result, REST is involved in numerous physiological and pathological processes. For example, REST was suggested to have a neuroprotective role in Alzheimer’s disease (AD) (Lu, Aron et al., 2014). In addition, longevity in humans was associated with REST upregulation and repression of neuronal excitation (Zullo, Drake et al., 2019).

Cochlear hair cells (HCs) and spiral ganglion neurons (SGNs) play essential roles in transmitting auditory signals. Inner hair cells (IHCs) convert mechanical stimuli into electrochemical signals transmitted to and conducted along auditory nerve fibers (D & Ricci, 2019, Fettiplace, 2017, Yu & Goodrich, 2014). In contrast, outer hair cells (OHCs) act as mechanical amplifiers that enhance weak sounds in the cochlea (Dallos, Wu et al., 2008, Liberman, Gao et al., 2002). SGNs are at the bottleneck between HCs and the brain, preserving sound information’s amplitude, frequency, and temporal features (Meyer & Moser, 2010, Taberner & Liberman, 2005). Since HCs and SGNs do not regenerate in the mature mammalian cochlea, their damage or dysfunction leads to permanent sensorineural hearing loss (Fujioka, Okano et al., 2015).

A recent study reported that the null deletion REST splice variant expressed in cochlear HCs is associated with DFNA27 hearing loss (Nakano, Kelly et al., 2018). Yet, the role of the full-length REST in hearing is mainly unknown. We generated a mouse model with REST conditionally knocked out in the cochlea. Results show that REST is expressed in cochlear HCs and SGNs of adult mice and that conditional deletion of REST in these cells resulted in progressive hearing loss. No detectable loss of cochlear HCs and SGNs was found at P1-P14 in development, but significant degeneration of OHCs and SGNs was detected at 3-months of age in mice. Examination of HCs and SGNs functions show that K_v_7.4 channel expression was upregulated, reducing SGNs excitability. Consistently, administration of the K_v_7.4 channel activator, fasudil, caused hearing loss in wild-type mice, while inhibition of the K_v_7.4 channel by XE991 rescued the hearing phenotype of *Rest* cKO mice. In sum, our results demonstrate that REST is essential for hearing and that REST deficiency causes upregulation of K_v_7.4 channels leading to dysfunction of SGNs and HCs and deafness in mice.

## Results

### REST is expressed in SGNs and HCs and is required for hearing

To examine the role of REST in hearing, we first examined the expression of REST in the cochlea of 1-month-old wild-type (WT) mice. REST was abundantly expressed in cochlear SGNs and HCs, as determined by immunofluorescence staining (**Figure 1A**) and single-cell RT-PCR (**Figure 1B**). We then investigated whether the specific deletion of *Rest* in the cochlea affects mouse hearing. Mice with homozygous intronic LoxP sites flanking exon 2 of *Rest* (*Rest^flox/flox^*) (Soldati, Bithell et al., 2012) were crossed with mice expressing the Cre recombinase transgene under the *Atoh 1*-specific promoter to produce *Rest* cochlear conditional knockout mice (*Rest* cKO) (**Figure 1C**). The genotypes of the pups were identified using PCR analysis (Fig. 1D).

**Figure 1.**
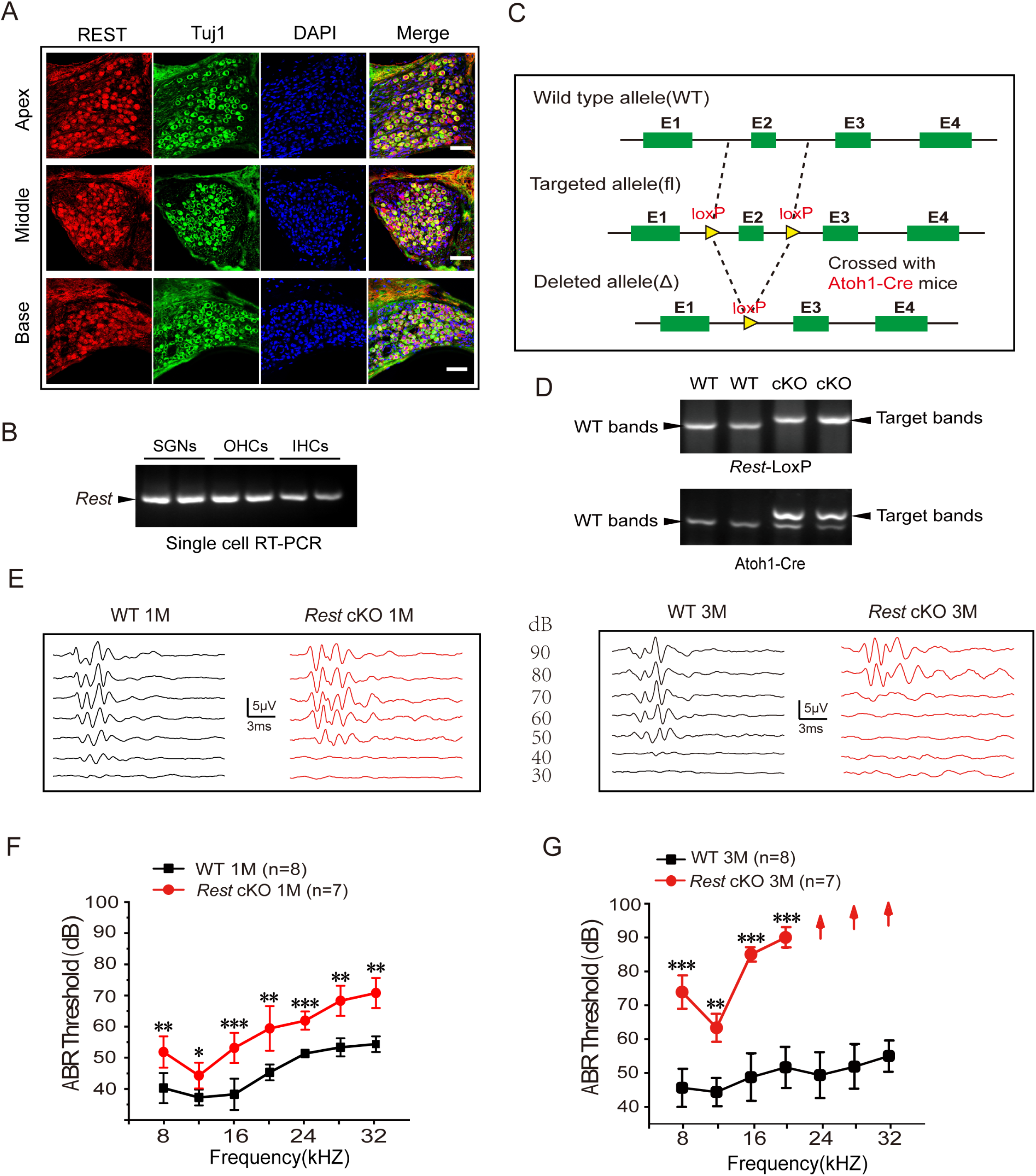
REST expression in the inner ear is essential for hearing. **A,** Expression of REST in SGNs from the apical, middle, and basal cochlea of WT mice. SGNs were stained using anti-REST (red) and anti-Tuj1, a neuron marker (green). The nuclei were stained with DAPI (blue). Scale bar: 20 μm**. B,** Single-cell RT-PCR analyses of *Rest* in SGNs, OHCs, and IHCs of WT mice**. C**, Schematic diagram of *Rest* conditional knockout (*Rest* cKO) generation process. Two LoxP sites were inserted into both alleles of *REST*, flanking the coding sequence of exon 2. Cre recombinase expression is activated by the specific *Atoh1* promoter in the cochlea and the exon-2 region between the homodromous LoxP sites, resulting in the loss of REST function cochlea. **D**, PCR genotyping of WT and *Rest* cKO mice using genomic DNA prepared from tail biopsies. *Rest* cKO mice were identified by two-step PCR, including first screening for mice containing the *Rest*-loxP target band and then further identifying for mice containing the *Atoh1*-cre target band. **E,** Representative ABR waveforms in response to clicking (90-30 dB) sound pressure levels in 1- and 3-month-old WT and *Rest* cKO mice. **F, G,** ABR threshold statistics of WT and *Rest* cKO mice at 1-month (**F**) and 3-months (**G**) of age in response to pure tone stimuli (8–32 kHz). Data are means±SEM, **p<0.01,***p<0.001.

The auditory function of 1-2-month-old WT and *Rest* cKO animals was evaluated by measuring ABR to pure tone stimuli. Both heterozygous and homozygous *Rest* knockout mice showed a significant increase in the ABR threshold (as compared to WT), indicating hearing impairment (**Figure 1-figure supplement 1**). There was no statistical difference in hearing decline between heterozygous and homozygous mice. Therefore, we used *Rest*-homozygous mice in subsequent experiments. We then further evaluated the hearing function of 1- and 3-month-old homozygous *Rest* cKO mice. Compared with age-matched WT mice, both 1- and 3-month-old *Rest* cKO mice displayed significantly threshold elevated across all frequencies (**Figures 1E-G**). Furthermore, the 3-month-old *Rest* cKO mice failed to elicit ABR waveforms at 24 kHz, 28 kHz, and 32 kHz, suggesting complete hearing loss in the high-frequency field (**Figure 1G**). These results indicate that *Rest* cKO mice display progressive hearing loss.

### Degeneration of SGNs and HCs is observed in 3-month-old but not 1-month-old *Rest* cKO mice

To investigate the cause of hearing loss induced by REST deficiency, we first examined the morphology of SGNs and HCs in *Rest* cKO mice. As shown in **Figures 2B and 2F**, immunofluorescence and HE staining revealed no apparent morphological abnormality or loss of SGNs and HCs in the cochlea’s apical, middle, and basal turns 1-month-old *Rest* cKO cochlea. In contrast, SGN and HC degeneration were observed in the cochlea of 3-month-old mice (**Figures 2C–E, G–L**). These data indicate that the loss of SGNs and HCs might be responsible for hearing loss in 3-month-old *Rest* cKO mice but not in 1-month-old mice.

**Figure 2.**
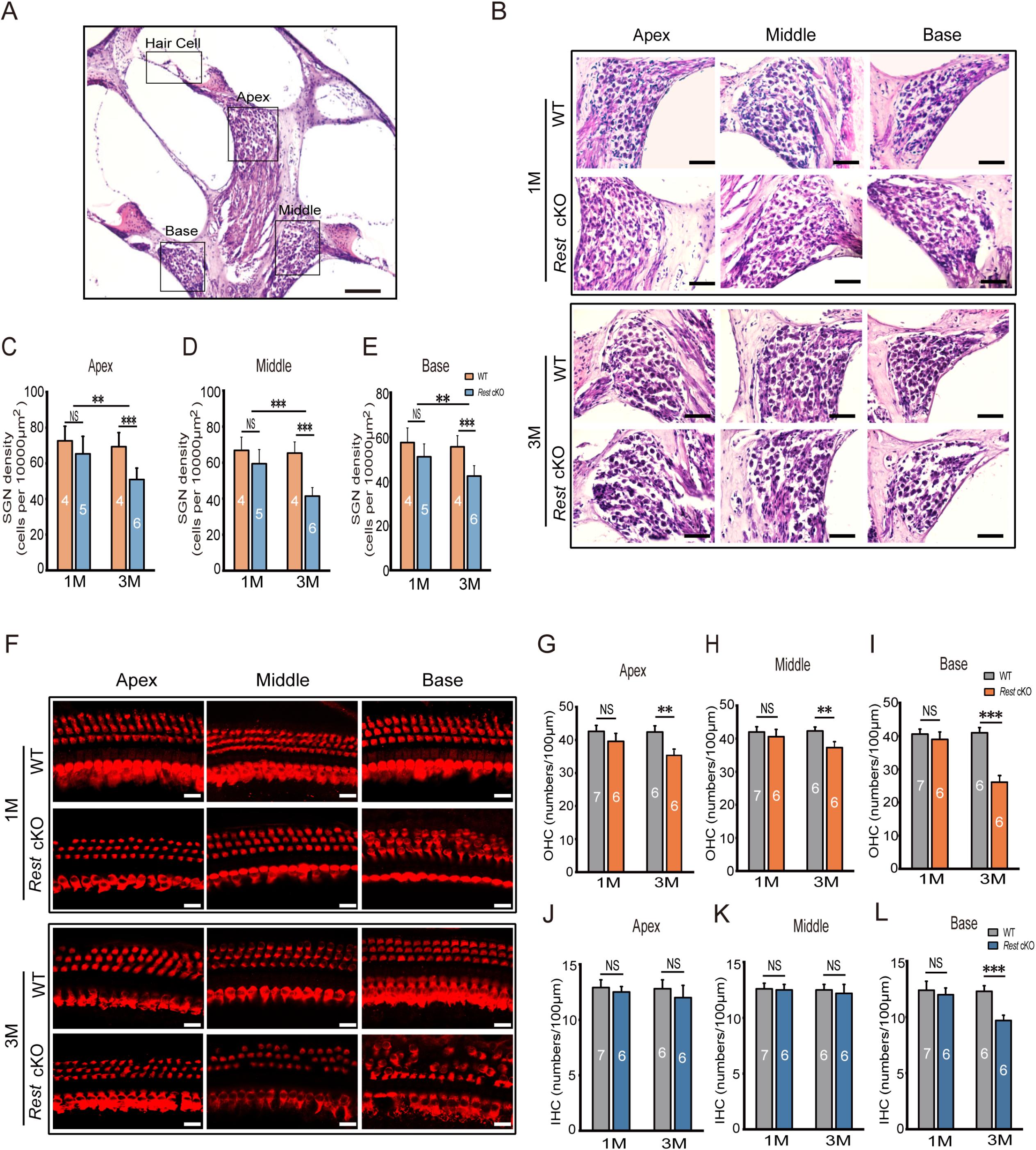
Degeneration of SGNs and HCs is observed in 3-month-old *Rest* cKO mice but not in 1-month-old mice. **A, B,** Morphometry changes in SGNs were observed in the apical, middle, and basal cochlea of 1- and 3-month-old WT and *Rest* cKO mice. Scale bar: 200 μm in A, and 50 μm in B. **C, D, E**, SGNs were counted by quantification in all three regions of the cochlea. **F,** Myo7a stained HCs from 1- and 3-month-old WT and *Rest* cKO mice. Scale bar: 20μm**. G–I,** Quantification of OHCs at the cochlea’s apical middle and basal regions in WT and *Rest* cKO mice (1M and 3M). **J–L,** Quantification data of IHCs in WT and *Rest* cKO mice. Data are means±SEM, **p<0.01,***p<0.001.

### The excitability of SGNs is decreased in *Rest* cKO mice

Given the lack of apparent indications of morphological alterations in SGNs and HCs in 1-month-old *Rest* cKO mice, we then explored whether dysfunction of SGNs or HCs resulted in the early onset of hearing impairment in these mice. We first examined the excitability of SGNs in *Rest* cKO and WT mice. The whole-cell current-clamp technique recorded action potentials (APs) in SGNs. A single spike was evoked by 0.4nA current injection in the apical SGNs of the cochlea in WT mice (**Figure 3A**). In contrast, 63.2% (1-month-old) and 57.6% (3-month-old) of SGNs from *Rest* cKO mice failed to produce APs, indicating reduced excitability of apical SGNs in *Rest* cKO mice (**Figure 3B**).

**Figure 3.**
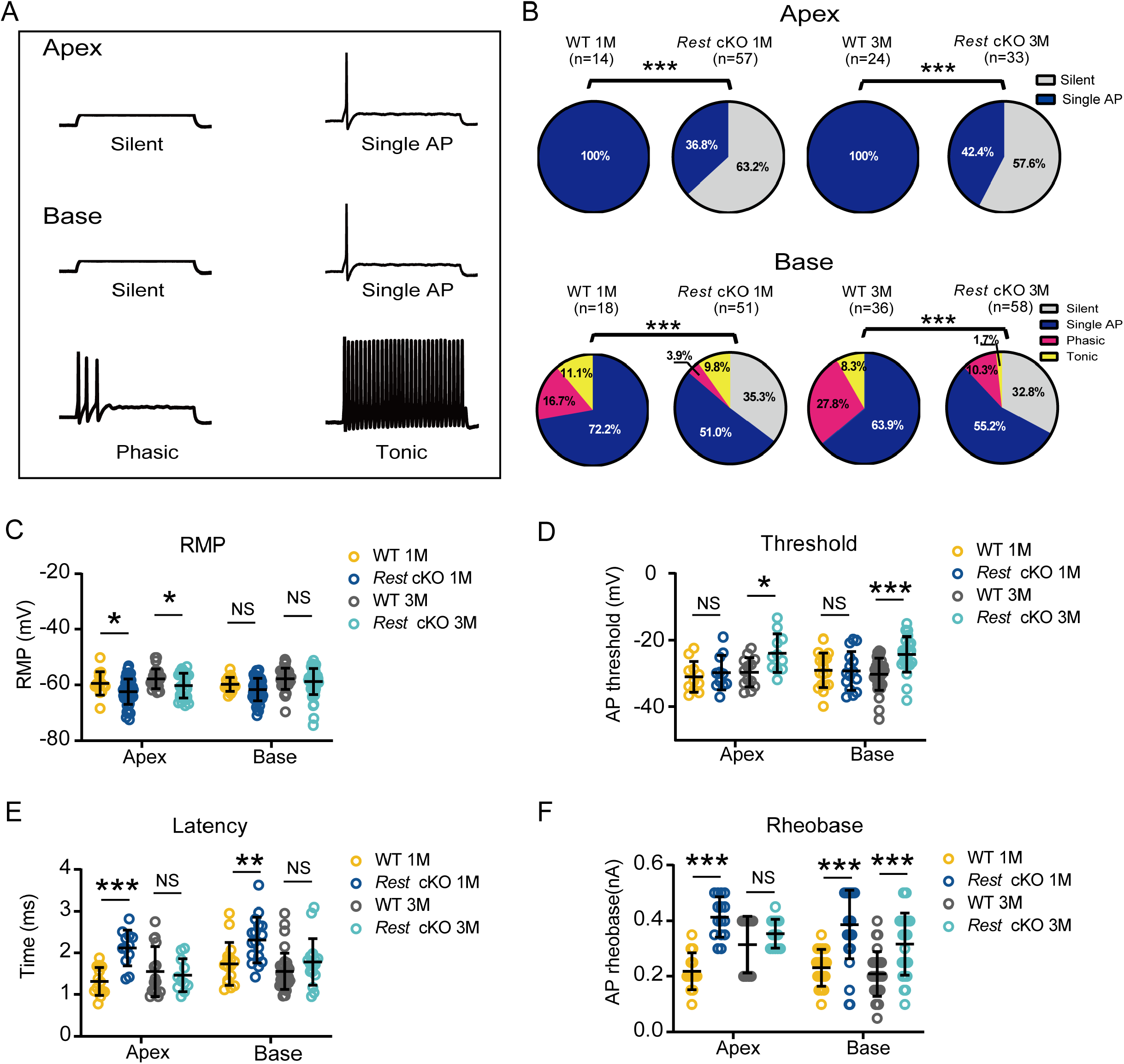
Reduced excitability of SGNs in *Rest* cKO mice. **A,** Representative traces show spike patterns of SGNs in the apical and basal cochlea, recorded with whole-cell patch clamps. Spikes were generated by 0.4 nA current injection. **B**, Pie charts illustrating the percentage abundance of the different spike patterns of SGNs in WT and *Rest* cKO mice. **C-F,** The resting membrane potentials (RMPs) (**C**), action potential thresholds (**D**), latencies (**E**), and rheobases (**F**) were recorded from apical and basal SGNs in WT and *Rest* cKO mice at 1 and 3 months. Data are means±SEM, *p < 0.05, **p<0.01,***p<0.001.

SGNs at the base of the cochlea in 1- and 3-month-old WT mice could generate multiple AP patterns, including single AP (1 spike), phasic (2–6 spikes), and tonic (>6 spikes) (**Figure 3A**). In contrast, the proportions of SGNs generating multiple spikes were decreased in *Rest* cKO mice, and there were a large number of basal SGNs failed to generate even a single AP. The proportions of these ‘silent’ neurons were 35.3% and 32.8% in 1- and 3-month-old *Rest* cKO mice, respectively. These recordings indicate that the excitability of SGNs at the base of the cochlea in *Rest* cKO mice was reduced (**Figure 3B**). The *Rest* cKO mice also displayed hyperpolarized resting membrane potentials (RMP), prolonged AP latencies, increased AP thresholds, and rheobase currents compared to WT mice (Figure 3C–F).

To confirm the effect of REST on the electrophysiological properties of SGNs, we recorded the excitability of SGNs in *Rest* cKO mice at P14 at the onset of hearing. Once again, SGN excitability was reduced at the apex and base (**Figure 3-figure supplement 1**). Thus, 27.8% of SGNs at the apex and 11.7% at the base failed to evoke APs. In addition, the RMPs of the SGNs at the base were hyperpolarized (**Figure 3-figure supplement 1**); significantly increased firing threshold and rheobase were recorded at the apex (**Figure 3-figure supplement 1**). Collectively, these data demonstrate that REST deficiency alters the electrophysiological properties of SGNs in mice, making these neurons less excitable.

### K_v_7.4 is upregulated in the inner ear of *Rest* cKO mice

Ion channels are the basis for controlling neuronal excitability. They are responsible for setting the resting membrane potential and maintaining aspects such as the duration, threshold, and firing frequency of action potentials. Therefore, we explored which ion channels are involved in the reduced excitability of SGNs induced by REST deficiency. Here, we focused on ion channels that regulate the excitability of mouse SGNs and contain RE-1 sites, such as K_v_4.3 and K_v_7.2-7.4(Mucha, Ooi et al., 2010, Reijntjes & Pyott, 2016, Uchida, Sasaki et al., 2010). We compared the changes in the expression of those channels in the cochlea of WT and *Rest* cKO mice. As shown in **Figures 4A and 4B**, K_v_7.4 channel mRNA and protein levels were significantly elevated in the cochlea of *Rest* cKO mice. Further immunofluorescence results showed increased expression of K_v_7.4 channel proteins in both HCs and SGNs (**Figures 4C–H**).

**Figure 4.**
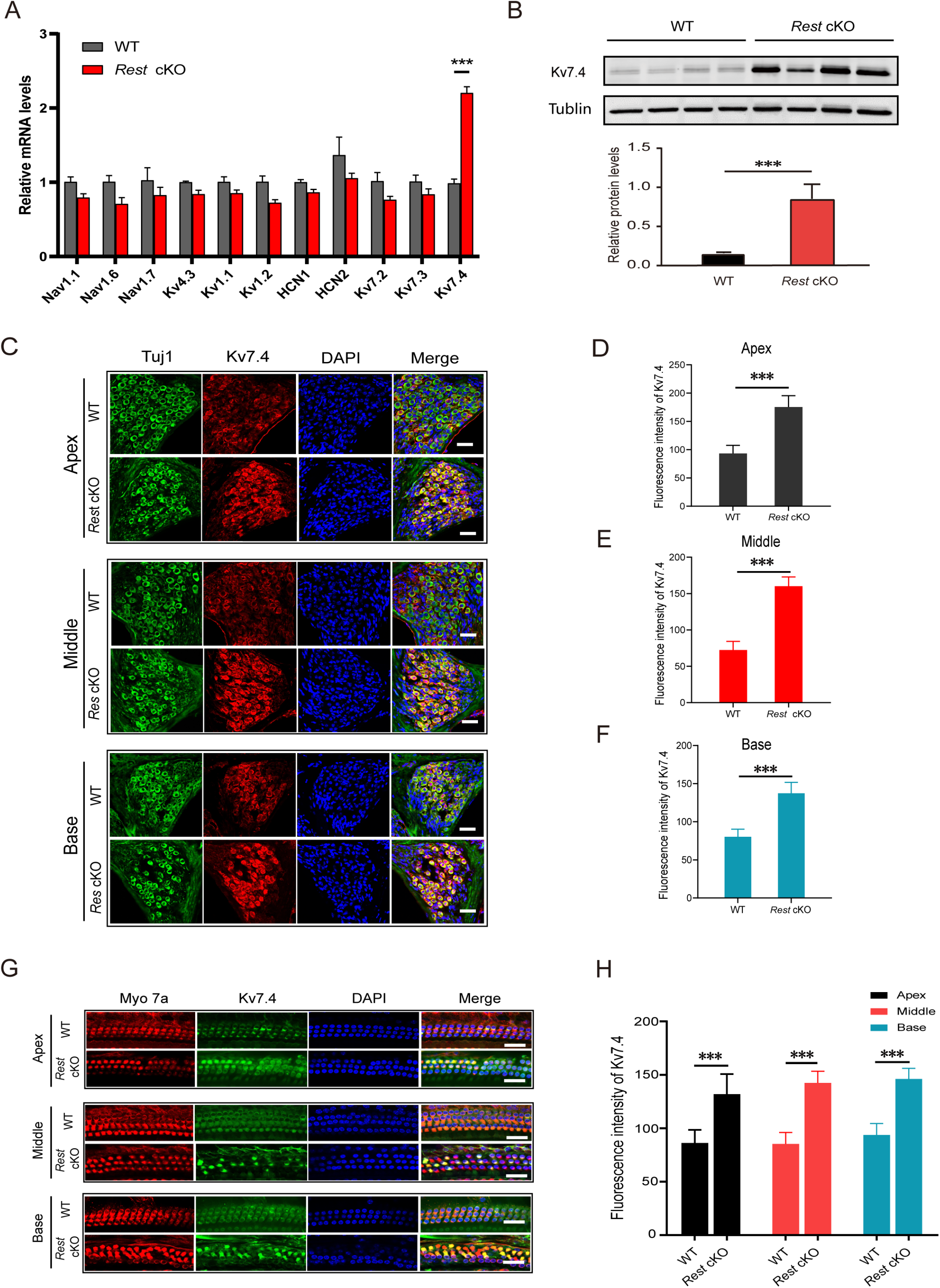
K_v_7.4 expression is increased in SGNs and HCs of *Rest* cKO mice. **A, B,** K_v_7.4 expression is significantly increased in the cochlea of *Rest* cKO mice at the mRNA and protein levels as determined by real-time PCR (**A**) and Western blotting (**B**). **C-F** Immunofluorescence shows increased K_v_7.4 (red) expression in the SGNs from apical, middle, and basal cochlea in *Rest* cKO mice. Scale bar is 20 μm in C, and G. **G, H,** K_v_7.4 expression is increased in HCs of *Rest* cKO mice. Data are means±SEM, ***p<0.001.

### Upregulation of K_v_7.4 caused a decrease in SGNs excitability in *Rest* cKO mice

K_v_7.4 channels belong to the K_v_7 potassium channel family and are important components of K_v_7 or ‘M-type’ neuronal K^+^ currents. To determine whether increased K_v_7.4 channel expression causes upregulation of channel function, we recorded K_v_7 channel currents in the SGNs of WT and *Rest* cKO mice. Using the whole-cell voltage-clamp technique, cells were held at −20 mV, then hyperpolarized to −60 mV, and the tail currents were recorded. The K_v_7 channel currents were identified as the current component, blocked by the K_v_7 channel blocker XE991 (3 μM) (**Figures 5A-C**). As shown in **Figures 5D** and E, K_v_7 channel current density was significantly increased in the SGNs at the apex and base of 1- and 3-month-old *Rest* cKO mice compared to WT mice. However, there was no significant upregulation of K_v_7 currents in the SGNs of P14 mice (**Figure 5-figure supplement 1**).

**Figure 5.**
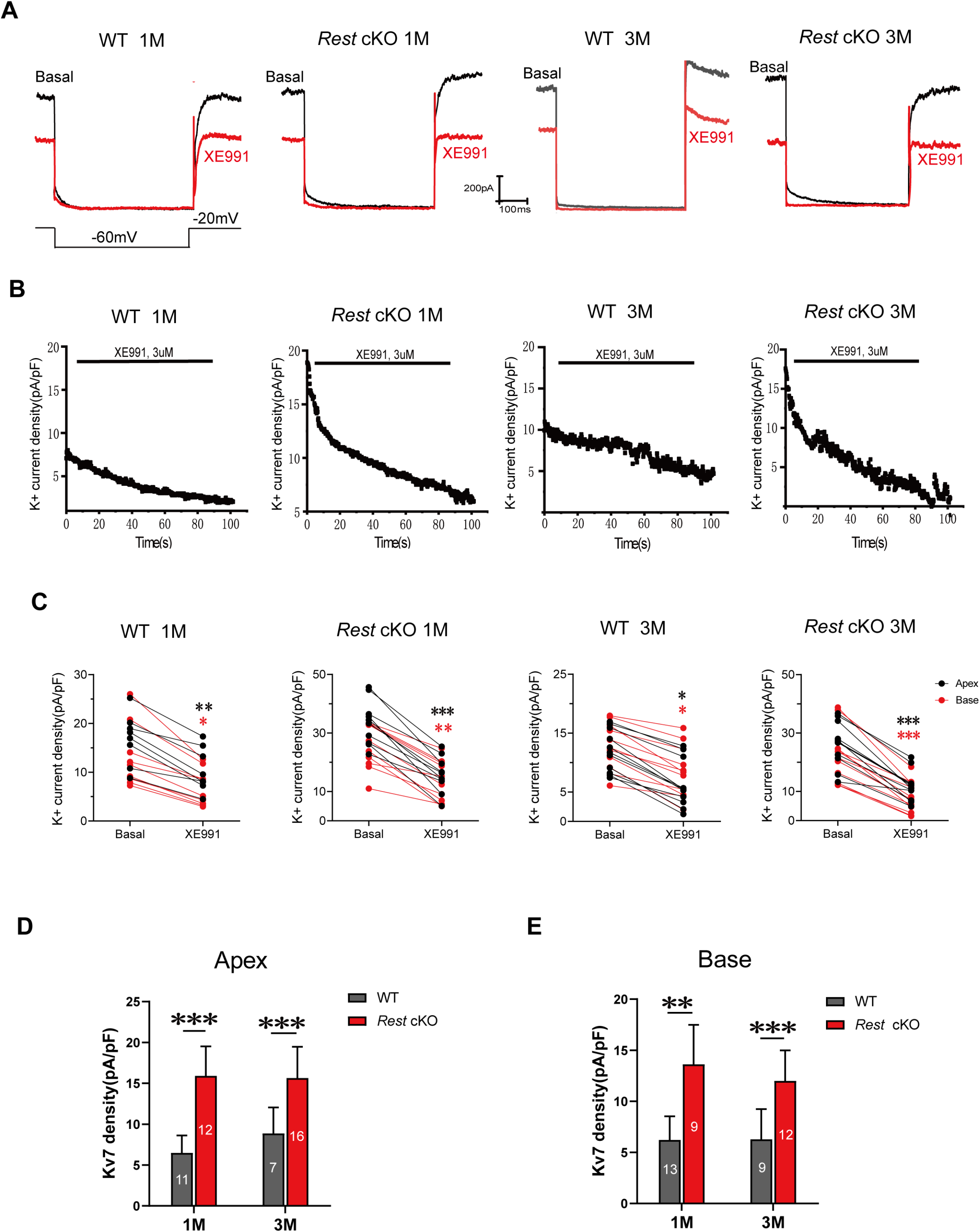
Increased K_v_7 currents in the SGNs of *Rest* cKO mice. **A**, Whole-cell K_v_7 currents were recorded in WT and *Rest* cKO SGNs. Representative K_v_7 currents were calculated as XE991-sensitive currents upon voltage step from −20 mV to −60 mV. **B**, Time course of outward K^+^ current inhibition by XE991 (3μM). **C**, Comparison of outward K^+^ current at −20 mV in individual SGNs in the absence (basal) and presence of XE991 (3μM). **D, E,** Summary data show that XE991-sensitive currents (K_v_7 current) density is significantly increased in the apical and basal SGNs of *Rest* cKO mice compared with WT mice. Data are means±SEM, *p < 0.05, **p<0.01, ***p<0.001.

K_v_7.2 and K_v_7.3 also contribute to K_v_7 channel currents in many neurons, and these are also expressed in SGNs (Jin, Liang et al., 2009). To determine whether K_v_7.2 and K_v_7.3 are involved in the upregulation of K_v_7 channel currents, we examined the mRNA levels of K_v_7.2 and K_v_7.3 in the cochlea of WT and *Rest* cKO mice using real-time PCR. As shown in **Figure 4A**, the expression levels of K_v_7.2 and K_v_7.3 were not significantly different between WT and *Rest* cKO mice. Further immunofluorescence staining confirmed that the expression of K_v_7.2 and K_v_7.3 channels was not changed considerably in the SGNs of *Rest* cKO mice (**Figure 5-figure supplement 2**). These data indicate that deletion of cochlear REST upregulates K_v_7.4 channels rather than K_v_7.2 and K_v_7.3 in SGNs. These results were unexpected as deletion of REST was shown to upregulate Kv7.2 in mouse sensory neurons(Zhang, Gigout et al., 2019) and may suggest that additional factors contribute to cell-type-specific regulation Kv7 channels by REST.

To further confirm the inhibitory effect of REST on K_v_7.4, we overexpressed REST in CHO cells stably expressing K_v_7.4 channels, and, indeed, this caused a significant decrease in K_v_7.4 current density (**Figure 5-figure supplement 3**). The Western blotting results confirmed that overexpression of REST significantly reduced the protein expression level of K_v_7.4 channels in CHO cells (**Figure 5-figure supplement 4**), indicating that REST can reduce K_v_7.4 currents by inhibiting K_v_7.4 protein expression.

### Upregulation of K_v_7.4 results in dysfunction of OHCs in *Rest* cKO mice

K_v_7.4 is the only K_v_7 family member reported to be expressed on OHCs so far, and it is necessary to maintain the electrophysiological properties of OHCs and their functions (Johnson, Beurg et al., 2011, Perez-Flores, Lee et al., 2020). We first recorded K_v_7 currents in the OHCs of P10-14 WT and *Rest* cKO mice. Whole-cell K^+^ currents were recorded from OHCs, and the XE991-sensitive currents denoted hereafter as K_v_7-mediated currents are shown in **Figures 6A-E**. As with SGNs, K_v_7 channel currents were significantly increased in the OHCs of *Rest* cKO mice (**Figure 6F**).

**Figure 6.**
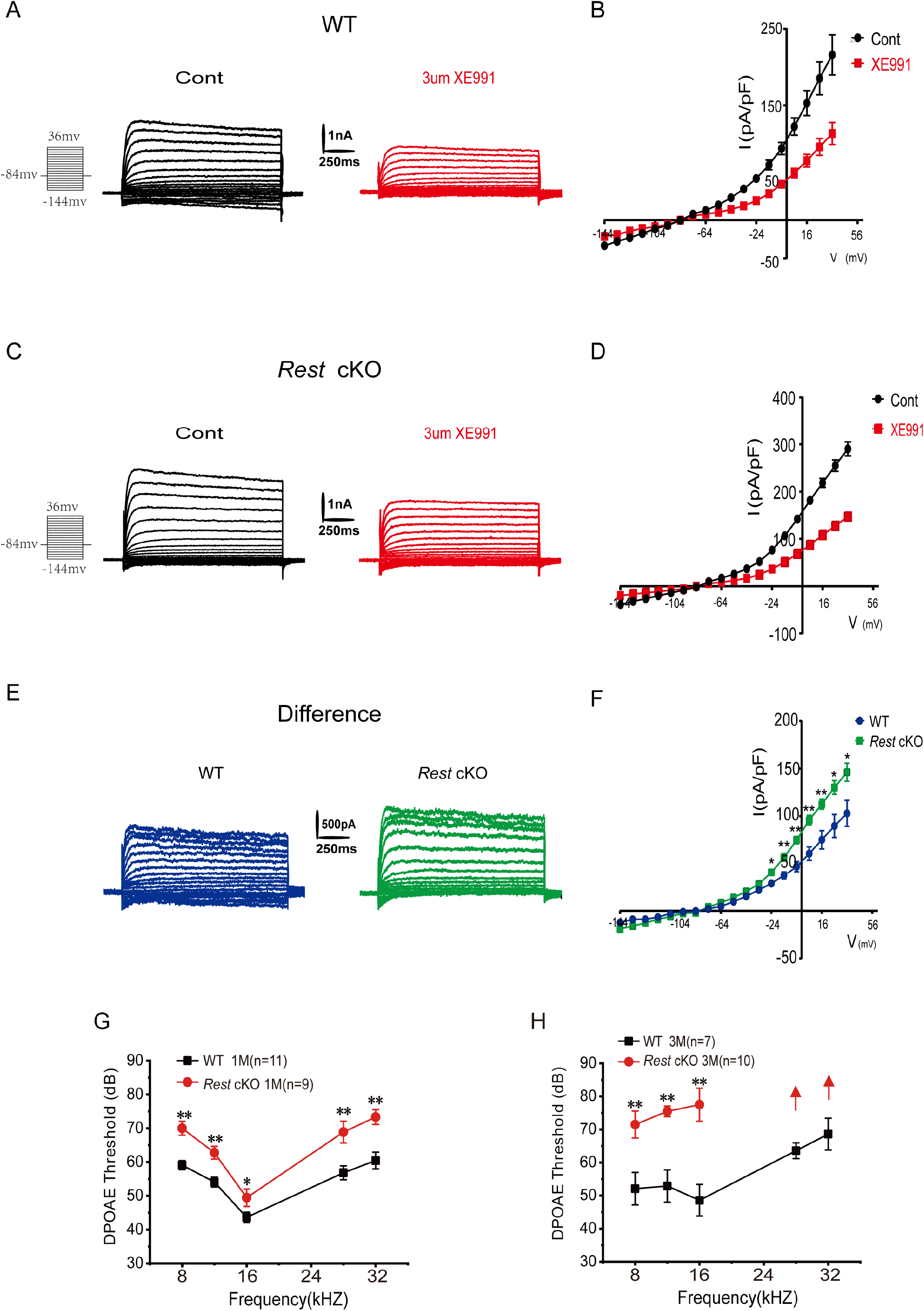
Increased K_v_7 currents in the OHCs of *Rest* cKO mice. **A,** Representative current traces recorded from the OHCs of P13 WT mice and the resulting traces after application of XE991. K^+^ currents were obtained from a holding potential of −84 mV and stepped up from-144 to +34 mV, with 10-mV increments (left panel). **B,** Current density-voltage curves are shown before and after XE991 (3μM) application in WT mice. **C**, Similar data was obtained from the OHCs of *Rest* cKO mice in the control condition, and after XE991 (3 μM), current traces were applied. **D,** Current density-voltage curves are shown before and after the application of XE991 in *Rest* cKO mice. **E,** The difference-current traces, which represent the XE991-sensitive currents. **F,** Difference current density-voltage curve obtained from OHCs in WT and *Rest* cKO mice. **G, H,** DPOAE threshold measurement of 1- and 3-month-old WT and *Rest* cKO mice. Data are means±SEM, *p < 0.05, **p<0.01.

Subsequently, we measured the distortion product otoacoustic emissions (DPOAEs). DPOAEs depend on active processes in the cochlea, which are linked to the normal function of OHCs (Dallos, 1992). DPOAEs detect low-level sounds produced by functional OHCs and emitted back to the ear canal. We demonstrated that DPOAE thresholds were significantly increased in *Rest* cKO mice (**Figures 6G-H**), indicating a decline in the function of OHCs; such an effect is consistent with the upregulation of K_v_7.4in OHCs.

### Pharmacological activation of K_v_7 channels impairs hearing in mice

To verify that upregulation of K_v_7.4 channel functions results in hearing loss *in vivo*, we administered fasudil (i.p., 10 mg/kg and 20 mg/kg), a K_v_7.4 channel-specific activator, to WT mice and evaluated their hearing function. ABR was measured before fasudil administration and 14 and 28 days after. As shown in **Figure 7A**, the hearing thresholds of mice in both 10 mg/kg and 20 mg/kg fasudil-treated groups were significantly higher on day 28, indicating hearing impairment in these mice (**Figure 7B**). We then determined whether the loss of SGNs or HCs induced hearing impairment. We demonstrated that all groups of mice displayed normal cochlear morphology with no loss of SGNs and HCs (**Figure 7-figure supplement 1**). These results suggest that fasudil treatment did not alter the morphology or number of HCs and SGNs.

**Figure 7.**
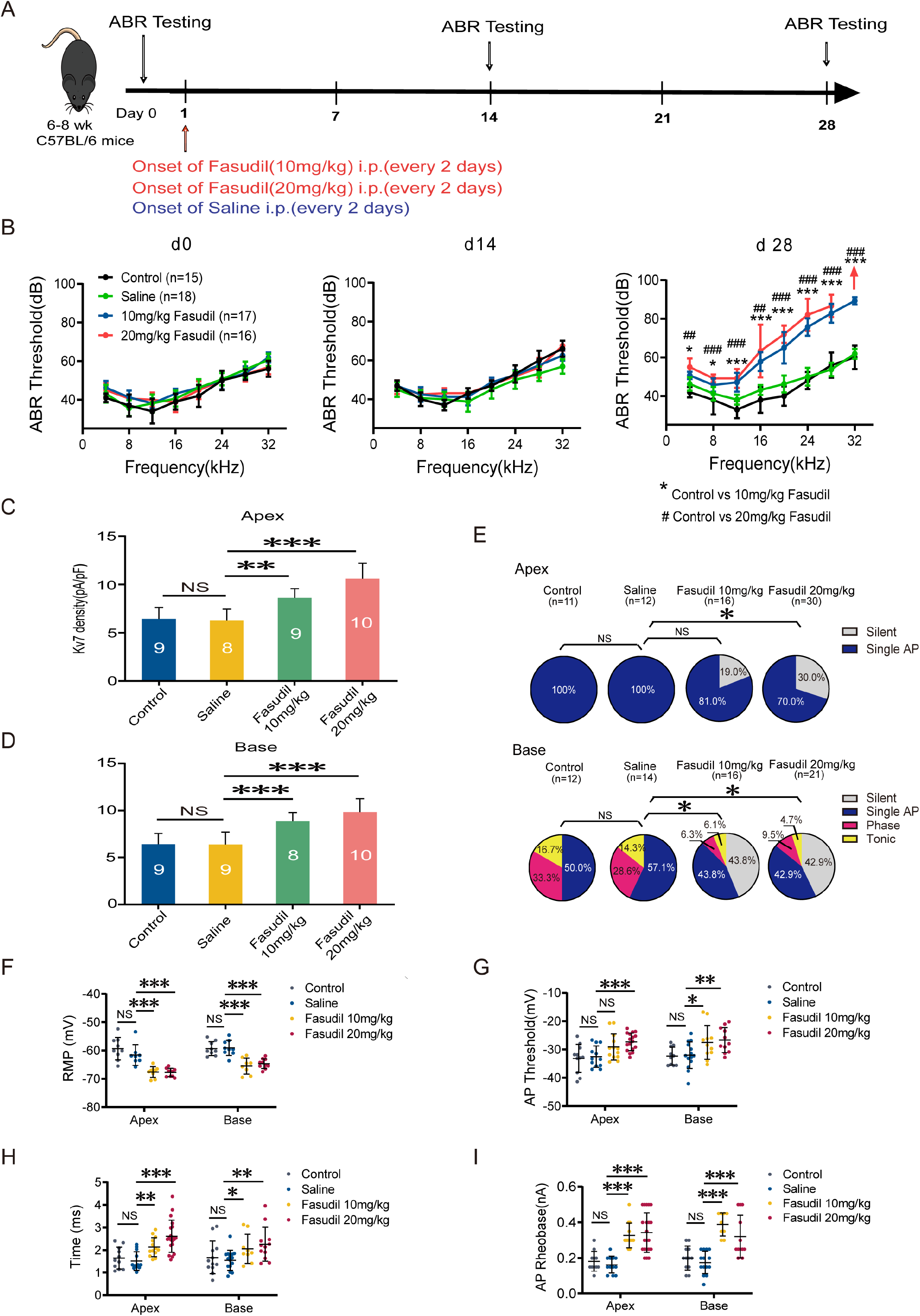
K_v_7.4 channel activation induced mice hearing impairments. **A**, Timeline for ABR testing and drug treatment. Fasudil was administered to 6–8 week-old WT mice intraperitoneally at 10 mg/kg or 20 mg/kg every other day for 28 days. **B,** ABR thresholds in control, saline, and fasudil (10 mg/kg, 20 mg/kg) treatment groups on days 0, 14, and 28. **C, D**, Summary data showing an increase in K_v_7 current density in the apical and basal SGNs of fasudil (10 mg/kg, 20 mg/kg)-treated mice. **E**, Summary of spike patterns recorded in SGNs from control, saline, and fasudil (10 mg/kg, 20 mg/kg)-treated mice. **F–I,** RMPs, action potential thresholds, latencies, and rheobases were recorded from the apical and basal SGNs of mice. Data are means±SEM, *p < 0.05, **p<0.01,***p<0.001.

We next assessed the effect of fasudil on cochlear cell function. We examined the effects of fasudil on K_v_7 currents in SGNs and the excitability of SGNs. K_v_7 current density on SGNs in both the apex and base was significantly increased in both fasudil-treated groups (**Figures 7C-D**). In addition, SGN excitability was dramatically reduced in the fasudil-treated group (**Figure 7E**), which was manifested with hyperpolarized RMPs, elevated AP thresholds, prolonged AP latencies, and elevated AP rheobases (**Figures 7F–I**).

### K_v_7 channel blocker XE991 rescues the hearing loss in *Rest* cKO mice

To confirm that the upregulation of K_v_7.4 channels caused by REST deficiency leads to deafness in mice, we investigated whether blocking K_v_7.4 channels rescues hearing loss in *Rest* cKO mice. We tested the hearing function of *Rest* cKO mice before and after the administration of XE991, a K_v_7 channel inhibitor (**Figure 8A**). The results showed that the ABR thresholds of *Rest* cKO mice were significantly reduced after treatment with 0.25 mg/kg XE991, indicating an improvement in their hearing (**Figures 8B-C**). Moreover, the ABR thresholds of these mice in the XE991 0.5 mg/kg treatment group were also significantly lower at low frequencies (4 to 8 kHz), indicating a partial improvement in their hearing (**Figures 8B-C**).

**Figure 8.**
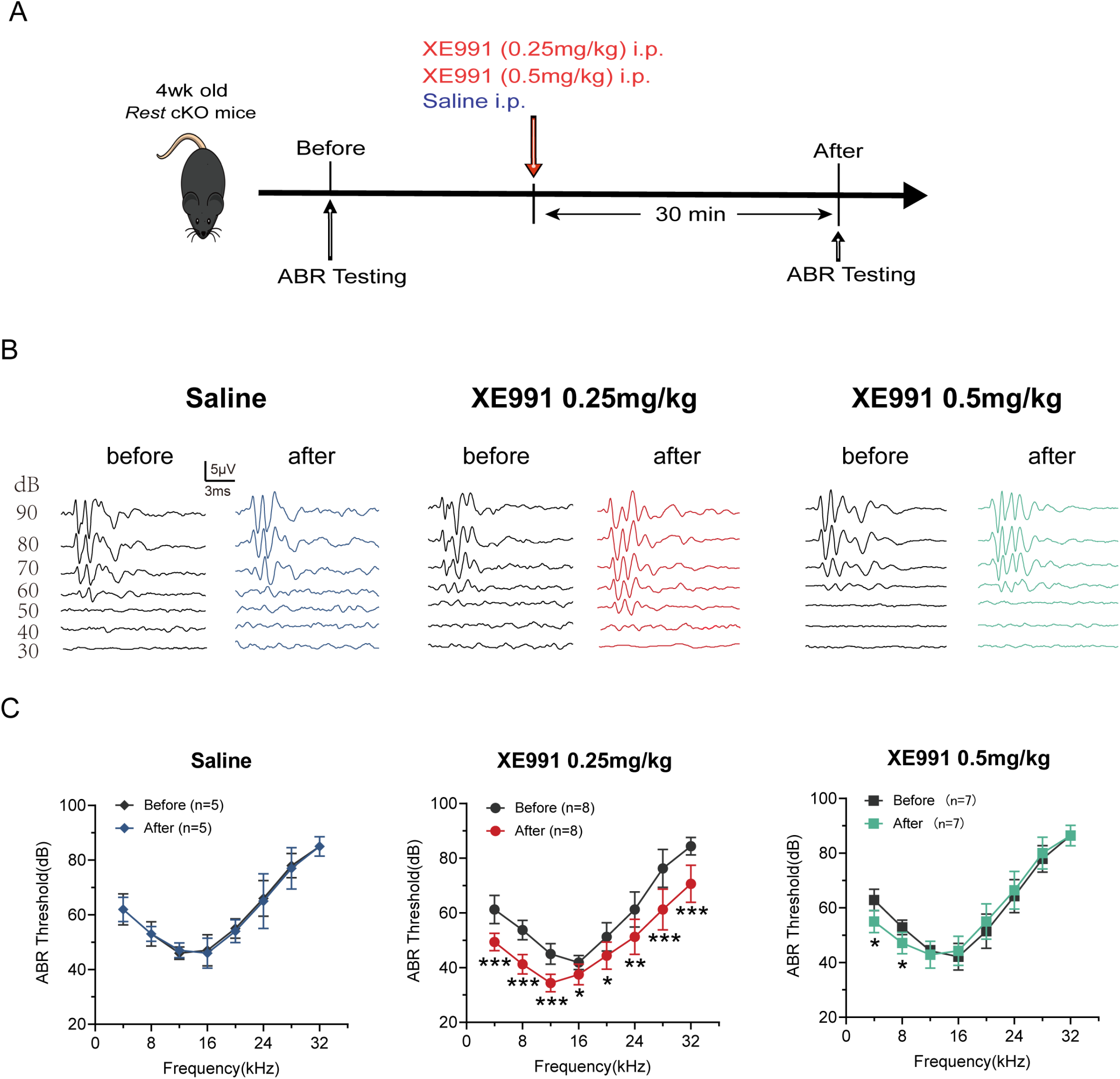
K_v_7 channel blocking rescues hearing of *Rest* cKO mice. **A**, Timeline for ABR testing and drug treatment. XE991 was administered to 1-month-old *Rest* cKO mice intraperitoneally at 0.25 mg/kg or 0.5 mg/kg. **B**, Representative ABR waveforms in response to clicking (90–30 dB) sound pressure levels in the mice of the saline and XE991-treated groups. **C**, ABR thresholds in the saline and XE991 (0.25, 0.5 mg/kg) treatment groups. Data are means±SEM, *p<0.05, **p<0.01, ***p<0.001.

Our results suggest that REST deficiency in the cochlea leads to increased expression of K_v_7.4 channels. Upregulation of K_v_7.4 function results in decreased excitability of cochlear SGNs and OHCs’ dysfunction and hearing loss in mice (**Figure 9**).

**Figure 9.**
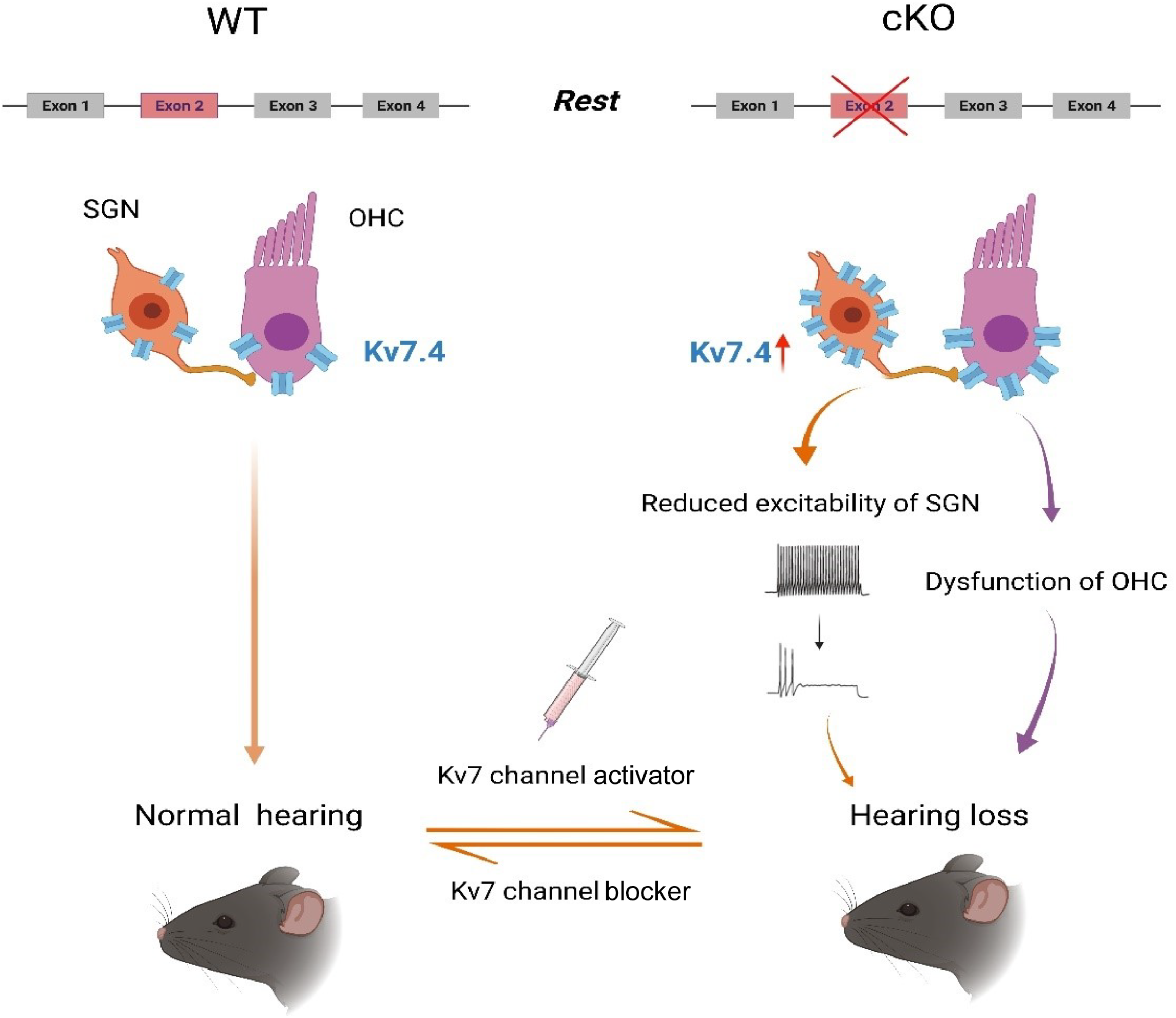
Model for the mechanisms of hearing loss in mice induced by REST deficiency through upregulation of K_v_7.4 channels. The illustration shows that REST deficiency in the cochlea leads to increased expression of K_v_7.4 channels and upregulation of channel function, which causes reduced excitability of SGNs and dysfunction of OHCs, ultimately leading to hearing loss in mice. Administration of K_v_7.4 channel activator, fasudil, recapitulated hearing loss in mice. In contrast, inhibition of the K_v_7.4 channel by XE991 rescued the auditory phenotype of *Rest* cKO mice. (The figure was created with BioRender.com)

## Discussion

REST, a transcriptional repressor, plays a dominant role in neural development and differentiation. Emerging evidence indicates that REST is an important player in the developed nervous system, including recognized roles in chronic pain development (Zhang et al., 2019) and aging (Lu et al., 2014, Zullo et al., 2019). However, the part of REST in hearing is still unknown. Here, we generated conditional deletion of *Rest* in HCs and SGNs. We found that specific deletion of *Rest* resulted in enhanced K_v_7.4 channels expression and increased K_v_7 currents in both cell types, leading to reduced SGN excitability and compromised HC function, with both effects resulting in progressive hearing loss in mice. Consistently, administration of the K_v_7.4 channel activator fasudil caused hearing loss in WT mice, while the K_v_7 channel inhibitor XE991 effectively rescued the hearing deficit of *Rest* cKO mice.

Several loss-of-function mutations of K_v_7.4 in humans and animal models were reported to cause gradual degeneration of HCs and SGNs, resulting in an autosomal dominant form of non-syndromic progressive hearing loss (DFNA2) (Coucke, Van Hauwe et al., 1999, Jentsch, Steinmeyer et al., 1990, Kharkovets, Dedek et al., 2006, Kim, Lv et al., 2011). Our findings that enhanced K_v_7.4 expression/current result in hearing loss reinforce a hypothesis whereby a certain range of basal K_v_7.4 functional expression is necessary for good hearing. In contrast, either reduction or increase outside this safety range causes cochlear dysfunction. Importantly, we identified the transcriptional suppressor REST as an essential regulator of hearing in mice.

The mammalian *Rest* gene is 20–30 kb in length and consists of four exons and three introns. *Rest* pre-mRNA can generate multiple splice variants with alternative splicing. Alternative splicing of *Rest* exon-4 was reported to reduce the repressive activity of REST, which is present in the HCs of P1–P7 mice (Nakano et al., 2018). Heterozygous deletion of this alternative exon resulted in the gain-of-function of REST, leading to HC degeneration and deafness, which is associated with DFNA27 hearing loss (Nakano et al., 2018), suggesting that *Rest* splice variants are necessary for HC development and hearing.

Here, we generated cochlea-specific *Rest* knockout mice using *Atoh1-Cre* mice crossed with *Rest* exon 2-flox mice to determine whether REST deficiency affects hearing. Because exon 2 is an essential exon encoding a functional REST protein, the knockout of exon-2 results in the loss of function of the full-length and all alternatively spliced REST variants. Hence, our results demonstrate that a total loss of functional REST in the cochlea caused hearing loss, suggesting that REST is vital for HC and SGN functions. REST was first localized in the chick cochlea by RT-PCR in 2002 (Roberson, Alosi et al., 2002). The REST splice variant was recently demonstrated in the immature HCs in mice (Nakano et al., 2018). This study found that REST is also expressed in adult HCs and mice’s immature and mature SGNs.

REST was initially identified as a transcriptional repressor regulating cell differentiation and development. We investigated whether REST deficiency-induced impairment of cochlear cell development accounts for possible developmental deficiencies in auditory pathways induced by Rest cKO. We examined the morphological changes in mice HCs and SGNs at P1-14 before the onset of hearing. Our results show no apparent morphological abnormalities, HCs, or SGNs loss in the cochlea of *Rest* cKO mice at P1-14 (**Figure 8-figure supplement 1**). This suggests that REST deficiency causes early onset of deafness independent of HC and SGN development. In contrast, 3-month-old *Rest* cKO mice showed losses in SGNs and HCs. Since REST regulates various genes involved in cell death (Lu et al., 2014), we reason such cell loss could be mediated by REST targets other than *Kcnq4*, especially given the fact that upregulation of K_v_7 channels is generally considered to be neuroprotective (Bierbower, Choveau et al., 2015, Gamper, Zaika et al., 2006). K_v_7.4 channel is a voltage-dependent potassium channel encoded by the *KCNQ4* gene. It belongs to the K_v_7 potassium channel family containing five K_v_7.1– K_v_7.5 (Jentsch, 2000, Jones, Gamper et al., 2021). The K_v_7 channels are widely distributed in neurons, the heart, vascular smooth muscle cells, and cochlea (Brueggemann, Mackie et al., 2014, Delmas & Brown, 2005, Jentsch, 2000, Kharkovets, Hardelin et al., 2000, Neyroud, Tesson et al., 1997). In the cochlea, K_v_7.4 is mainly expressed in the OHCs, IHCs, and SGNs (Beisel, Rocha-Sanchez et al., 2005, Dierich, Altoe et al., 2020, Kharkovets et al., 2000). When expressed in heterologous expression systems, K_v_7.4 channels slowly activate and deactivate non-inactivating currents (i.e., as shown in Figure 5-figure supplement 3). We noted that the XE991-sensitive currents recorded from OHCs had faster kinetics than expected from heterologously expressed K_v_7.4 homo-tetramers (**Figure 6E**). Hence the OHCs may contain endogenous factors or modulators that alter the K_v_7.4 channel kinetics. On the other hand, currents recorded from SGNs had an ‘M-like’ slow-kinetics component, especially in neurons from *Rest* cKO (**Figure 5-figure supplement 3**). However, loss of function mutations in *KCNQ4* cause DFNA2 sensorineural deafness (Coucke et al., 1999, Kharkovets et al., 2006). Our results demonstrate that upregulation of K_v_7.4 channels also leads to hearing loss in mice by silencing SGNs and inducing dysfunction in HCs.

Here we did not directly measure the binding of REST to *Kcnq4,* but direct binding has been reported before (Iannotti, Barrese et al., 2013). REST regulates the expression of a variety of ion channel genes, including genes coding for HCN channels (Kuwahara, 2013, McClelland, Flynn et al., 2011), Nav1.2 channels (Armisen, Fuentes et al., 2002, Pozzi, Lignani et al., 2013), K_v_7.2/K_v_7.3 potassium channels (Mucha et al., 2010, Rose, Ooi et al., 2011), and T-type calcium channels (van Loo, Schaub et al., 2012). Although some ion channel-related target genes of REST were also expressed in the cochlea, our results showed that REST deletion only significantly upregulated K_v_7.4 channels in SGNs and HCs (amongst genes tested here, see **Figure 4A**). The reasons why REST deletion specifically upregulates *Kcnq4* but not other RE1-containing genes remain to be investigated. However, it is worth noting that different RE1 sites bind REST with different affinities(Ooi & Wood, 2007); hence, at least some of potentially REST-targeted genes may not be tonically suppressed (and, thus, not affected by REST deletion). In addition, REST could suppress additional regulators of other genes. Therefore, the outcome of REST deletion may not necessarily affect the expression of all RE1-containing genes in the same way. Moreover, the pattern of involved genes may be cell- and tissue-specific.

To verify that upregulation of K_v_7.4 channels can cause hearing loss, we administered the K_v_7.4 channel opener, fasudil. Fasudil was initially described as a Rho-associated protein kinase (ROCK) inhibitor (Shi & Wei, 2013). However, recently Li and colleagues (Li, Sun et al., 2017) found that fasudil selectively increased K_v_7.4 channel currents in dopaminergic (DA) neurons in the ventral tegmental area (VTA), as well as heterologous K_v_7.4 in the expressed system. Later, the same group reported that fasudil selectively activated K_v_7.4 channels in vascular smooth muscles (Zhang, An et al., 2016). It was further demonstrated that fasudil has a selective opening effect on K_v_7.4 potassium channels without significantly affecting other members of the K_v_7 channel family (Zhang et al., 2016). Here, we found increased K_v_7 potassium channel currents in SGNs in mice 28 days after recurrent (every 2 days) fasudil administration, an effect accompanied by deafness. It has to be noted that K_v_7.4 channels are also expressed in neurons of the dorsal nucleus of the cochlea and the hypothalamus nucleus in the auditory conduction pathway (Kharkovets et al., 2000). Thus, fasudil could influence hearing by affecting K_v_7.4 channels in these neurons. Hence, the exact site of action of systemic fasudil on hearing requires further elucidation.

In conclusion, our findings revealed that a critical homeostatic level of functional expression of the K_v_7.4 channel is required for proper auditory functions. REST is necessary for hearing through regulating such functional expression of K_v_7.4. Our study contributes to an understanding of the essential functions of REST in the auditory system. It provides new insight for potential future treatments of hearing disorders associated with the K_v_7.4 channel.

## Materials and Methods

### Animals

All experimental animal protocols were performed following the Animal Care and Ethical Committee of Hebei Medical University (Shijiazhuang, China). The mice were bred and housed under a 12:12 light-dark cycle. To generate *Atoh1-Cre: Rest^flox/flox^* conditional knockout mice (*Rest* cKO), *Rest^flox/flox^* mice (Soldati et al., 2012) were crossed with *Atoh1-Cre* mice (Jackson Laboratories, Bar Harbor, ME, stock No. 011104). Genotyping of the mouse tails and cochlear tissues was performed using the primers listed in Table1.

### Auditory Brainstem Responses (ABRs) and Distortion product otoacoustic emissions (DPOAE)

ABR measurements were performed as previously described (Shen, Liu et al., 2018). Briefly, mice were anesthetized with ketamine (100 mg/kg) and xylazine (10 mg/kg). Three electrodes were inserted subcutaneously at the vertex of the head (reference), ipsilateral mastoid (recording), and contralateral rear leg of the mice (ground). At 8, 12, 16, 20, 24, 28, and 32 kHz, click or tone stimuli were emitted from a 20 dB with an attenuation interval of 5 dB. ABRs were measured using a System III workstation (Tucker Davis Technologies, Alachua, FL, USA) in an IAC BioSigRP Soundbooth (GM Instruments, Irvine, UK). The hearing threshold was defined as the lowest sound intensity required to generate a reproducible ABR waveform.

The DPOAE at 2f1-f2 were elicited from test mice using BioSig-RP software and a TDT system (Tucker-Davis Technologies). Five frequency points of 4, 8, 16, 28, and 32 kHz were selected to measure the 2f1-f2(f2/f1=1.2) to predict the auditory thresholds. Hearing thresholds were defined as the averaged signal for each identified frequency tested and compared with the corresponding frequency in the controls.

### Cell culture

SGNs were isolated from the mouse inner ears following a detailed procedure outlined in a previous study (Lv, Wei et al., 2010). Mice at various ages were humanely killed, the temporal bones were removed in a solution containing minimum essential medium (MEM) with Hank’s Balanced Salt Solution (HBSS) (Invitrogen) supplemented with 0.2 g/L kynurenic acid, 10 mM MgCl_2_, 2% fetal bovine serum (FBS; v/v), and glucose (6 g/L). The SGN tissue was then dissected and split into two segments: the apex and base across the modiolar axis. The two-segment tissue was digested separately in an enzyme mixture containing collagenase type I (1 mg/ml) and DNase (1 mg/ml) at 37 °C for 15 min. After gentle trituration and centrifugation (2,000 rpm for 5 min) in 0.45 M sucrose, the cell pellets were reconstituted in 900 μl of culture media (Neurobasal^TM^ A, supplemented with 2% B27 (v/v), 0.5 mM L-glutamine, 100 units/ml penicillin; Invitrogen). The suspension containing the SGNs was filtered through a 40-μm cell filter and planted on glass coverslips pretreated with 0.5 mg/ml poly D-lysine (Sigma-Aldrich) and 1 mg/ml laminin (Sigma-Aldrich). SGNs were kept in culture for 24-48h before electrophysiological recordings to allow for the detachment of Schwann cells from the soma.

A Chinese hamster ovary (CHO) cell line stably expressing K_v_7.4 channels (Inovogen Tech, China) was maintained in MEM with 10% fetal bovine serum (FBS), a 1% penicillin-streptomycin mixture (Gibco), and 20μg/mL puromycin at 37 °C with 5% CO_2_. CHO cells were cultured in MEM supplemented with 10% FBS without antibiotics for 12–24 h before transfection. The pIRES2-REST-EGFP (Sangon Biotech, Shanghai, China) or the pIRES2-EGFP (Sangon Biotech, Shanghai, China) plasmid constructs were transiently transfected into CHO cells alone or in combination, at a total amount of 600ng/well, using Lipofectamine 3000 (Invitrogen) according to the manufacturer’s instructions.

### Electrophysiology

Whole-cell voltage-clamp recordings from SGNs were performed at room temperature using an Axopatch 700 B amplifier (Molecular Devices, Sunnyvale, CA, USA). Signals were filtered at 2 kHz with a low-pass Bessel filter and digitized at ≥20 kHz using a 12-bit acquisition system, Digidata 1332 (Axon Instruments), and pClamp 10.7 software (Molecular Devices, Sunnyvale, CA, USA). Patch pipettes were pulled from borosilicate glass capillaries and heat polished to a tip resistance of 3–4 MΩ. The pipette solution contained (in mM): 112 KCl, 2 MgCl_2_, 0.1 CaCl_2_, 10 HEPES, 1 EGTA, 5 K_2_ATP, and 0.5 Na-GTP, adjusted to a pH of 7.35 with potassium hydroxide (KOH). The bath solution contained 130 mM NaCl, 5, 1 mM MgCl_2_, 2 mM CaCl_2_, 10 mM HEPES, and 10 mM glucose, adjusted to a pH of 7.4 with NaOH. The magnitude of the K_v_7 channel currents in the SGNs was measured from the deactivation current when stepping the membrane voltage from −20 mV to − 60 mV. It was calculated as the difference between the current amplitude at 10-ms into the voltage pulse and that at the end of the pulse. XE991 was purchased from Tocris Bioscience, and all other chemicals were purchased from Sigma.

Whole-cell current-clamp recordings from SGNs were conducted using the same pipette solution used in the voltage-clamp experiment. To measure the K_v_7.4 channel currents in CHO cells stably expressing K_v_ 7.4 channels, pipettes were filled with internal solutions containing (in mM): 145 KCl, 1 MgCl_2_, 10 HEPES, 5 EGTA, 5 K_2_ATP, adjusted to a pH of 7.35 with KOH. The external solution contained 140 mM NaCl, 3 KCl, 1.5 MgCl_2_, 2 CaCl_2_, 10 HEPES, and 10 mM glucose and was adjusted to 7.4 pH with NaOH. K_v_7.4 channel current traces were generated with 2.5 s depolarizing voltage steps from a holding potential of −80 mV and stepped to a maximal value of +40 mV in 10 mV increments. Currents were measured with capacitance and series resistance compensation (nominally 70–90%).

To record K^+^ currents in outer hair cells (OHCs), apical cochlear turns of P10– P14 mice were dissected in a solution containing (in mM): 144.6 NaCl, 5.5KCl, 1 MgCl_2_, 0.1 CaCl_2_, 0.5 MgSO_4_, 10.2 HEPES, and 3.5 L-Glutamine, adjusted to a pH of 7.2 with NaOH. The pipette solution contained (in mM): 142 KCl, 3.5 MgCl_2_, 1 EGTA, 2.5 MgATP, 0.1 CaCl_2_, 5 HEPES, adjusted to a pH of 7.4 with KOH. The external solution contained (in mM): 145 NaCl, 5.8 KCl, 0.9 MgCl_2_, 1.3 CaCl_2_, 0.7 NaH_2_PO_4_, 10 HEPES, and 5.6 D-glucose, adjusted to a pH of 7.4 with NaOH. To record K^+^ currents in OHCs in a whole-cell configuration, OHCs were held at −84 mV with step voltages ranging from −144 mV to 34 mV, with 10 mV increments. Recordings were performed at room temperature using Axon patch 700 B (Molecular Equipment, USA) amplifiers. Data acquisition was controlled by Clampex10.7 (Molecular equipment, USA) and a Digidata 1440A A-D converter (Molecular equipment, USA).

### Whole-mount preparation of SGNs and HCs

The mice were anesthetized, and the cochleae were fixed in 4% paraformaldehyde (PFA) overnight at 4 °C and then decalcified in 10% EDTA (#798681, Sigma, Darmstadt, Germany) in PBS for 2–3 days at 4 °C. The EDTA solution was changed daily. Decalcified cochleae were processed using a sucrose gradient and embedded in an OCT compound (Tissue-Tek) for cryosectioning. The specimens were sliced into 10-μm sections for study.

For HC preparation, the SGNs, Reissner’s membrane, and the most basal cochlear segments were removed after fixation with 4% PFA for 2 h, followed by decalcification in 10% EDTA for 3-days at 4°C. HC preparations were used for immunofluorescence staining.

### Cell count

Cochlear sections were stained with hematoxylin and eosin (HE) to evaluate SGN morphometry and density. Rosenthal’s canal was divided into the apex, middle, and base regions. SGN density from these three regions was measured using Image-Pro Plus 5.1. Five to six sections per cochlea per animal were analyzed, and six mice were used in each group. Cells displaying Myo7A labeling were quantified as the number of positive cells per 100 μm of basilar membrane length from the apex, middle, and base for HC counts.

### Immunofluorescence staining

Specimens were simultaneously permeabilized and blocked with 10% goat serum in 0.1% TritonX-100 and 0.1% BSA for 30 min at 37 °C and then labeled with primary antibodies overnight at 4°C. The primary antibodies used in this study were mouse anti-K_v_7.4 antibodies (#ab84820, Abcam, Cambridge, UK; 1:100), rabbit anti-REST antibodies (#ab21635, Abcam, Cambridge, UK; 1:200), mouse anti-Tuj1 (#801202, Biolegend, CA, USA; 1:100), rabbit anti-Myo7A antibodies (#25-6790, Proteus BioSciences, CA, USA; 1:50), and Phalloidin-iFluor 555(#ab176756, Abcam, Cambridge, UK). For detection, the specimens were incubated with Alexa 488-conjugated and Alexa 568-conjugated secondary antibodies (#115-545-003, 115-165-003, 111-545-003,111-165-003, Jackson Immuno Research, PA, USA; 1:400) for 2-h at room temperature and then stained with 4,6-diamidino-2-phenylindole dihydrochloride (DAPI, #D9542, Sigma, Darmstadt, Germany) for 10 min at room temperature. After washing with PBS, ProLong™ Gold Antifade Mountant (# P36934, Invitrogen, CA, USA) was used to mount the samples. Sections were visualized using a confocal fluorescent microscope (Leica Microsystems, Wetzlar, Germany). Images were analyzed with Leica LAS AF Lite and processed using ImageJ and Photoshop CS5 (Adobe, San Jose, CA).

### Single-cell RT-PCR

Cochleas were dissected from adult WT mice. SGNs or HCs were aspirated into a patch pipette using a patch-clamp system under the microscope. The electrode tip was then quickly broken into an RNase-free PCR tube containing 1 µL of oligo dT primers (50 µM), 1 µL of dNTP mixture (10 mM), 2 µL of RNase free water. The mixture was incubated in the water bath at 65℃ for 5 min and then placed on ice for 1 min. Single-strand cDNA was synthesized from cellular mRNA by supplementing with 1 µL of PrimeScript II RTase (200 U/µl), 0.5 µL of RNase Inhibitor (40 U/µl), 2 µL of 5×PrimeScript II buffer, and 1.5 µL of RNase Free H_2_O and then incubating the mixture at 55℃ for 50 min followed by 85℃ for 5 min. The first strand synthesis was performed at 95℃ for 5 min, followed by 35 cycles (95 ℃ for 50 s, 59 ℃ for 50 s, 72 ℃ for 50 s) and final elongation at 72 ℃ for 10 min by adding “outer” primers (Suppl. Table 3). Then, 2 µL of the first PCR products were used as the DNA template to amplify the second round PCR reaction by adding short-chain primer pairs, followed by 35 cycles (95 ℃ for 50 s, 59 ℃ for 50 s, 72 ℃ for 50 s) and the last elongation at 72 ℃ for 10 min. The second PCR product was identified by 2% agarose gels electrophoresis. The reverse transcription kit was purchased from Takara-Clontech (6210A, Invitrogen, USA), and the first second PCR kits were obtained from Promega (M7122, Madison, USA).

### Western blotting

Cochlear tissues (five mice per sample) were homogenized, and cells were lysed in a RIPA solution containing 0.1% SDS, 50 mM Tris (pH 7.4), 1mM EDTA (pH 8.0), 1% sodium deoxycholate, 1% TritonX-100, and 200 μM of phenylmethanesulfonyl fluoride (PMSF). The lysis mixture was centrifuged at 13800g for 30 min at 4°C. The supernatant was removed, and the protein concentration was determined using a BCA Protein Assay Kit (#23235, Thermo, CA, USA). Equal amounts of protein (40 μg) were resolved by SDS-PAGE and transferred to a PVDF membrane (#IPVH00010, Millipore, MA, USA). Membranes were blocked with 1×TBST containing 5% dry milk for 2-hr at room temperature and probed with the following antibodies: anti-K_v_7.4 antibodies (#ab84820, Abcam, Cambridge, United Kingdom; 1:500), anti-REST antibodies (#ab21635, Abcam, Cambridge, United Kingdom; 1:1000), or anti-tubulin beta-III antibodies (#MMS-435P, Biolegend, CA, USA; 1:10000) overnight at 4°C. Then, fluorescent secondary conjugated IgG (#IRDye^®^ 800CW goat anti-rabbit IgG (H + L); #IRDye^®^ 800CW goat anti-mouse IgG (H + L), LI-COR Biosciences, NE, USA; 1:10000) was applied for 90 min at room temperature. All Western blots were quantified using Image Studio software (IS, LI-COR Biosciences, NE, USA).

### Quantitative Real-Time PCR

Total RNA from mouse cochleae was extracted using an RNA Extraction Kit according to the manufacturer’s instructions (#9767, TaKaRa, Japan). The extracted RNA concentration was measured using a Nanodrop 2000 spectrophotometer (#ND-LITE, Thermo, DE, USA), and 1000 ng of total RNA was reversed transcribed using PrimeScript reverse transcriptase (#RR036Q, TaKaRa, China). The synthesized cDNA was used as a template to perform quantitative real-time (qRT)-PCR with TB Green^®^ Premix Ex Taq™ (#RR420Q, TaKaRa, China) in a Bio-Rad Real-Time PCR System (Bio-Rad Laboratories, CA, USA). Glyceraldehyde 3-phosphate dehydrogenase (GAPDH) was used as an internal control to normalize the relative mRNA abundance of each cDNA. The relative mRNA expression levels were calculated using the standard formula 2^-ΔΔCt^. All the primers used in this study were designed using Primer Premier 6.25 (PREMIER Biosoft International, CA, USA) and synthesized by Sangon Biotech (Shanghai, China). The primer sequences are shown in Table 2.

### In vivo drug administration

For fasudil administration: six- to eight-week-old C57 mice were divided into four groups: control, saline, fasudil 10 mg/kg, and fasudil 20 mg/kg. Fasudil was obtained from the National Institute for the Control of Pharmaceutical and Biological Products (Beijing, China), dissolved in 0.9% saline and administered intraperitoneal injection every two days for 28 days. Control mice were injected intraperitoneally with the same volume of saline. ABR was recorded before and after treatment with the drug and on days 14 and 28 after treatment to assess hearing function. The cochlea was also removed on day 28 to evaluate morphological changes of the HCs and SGNs and the functional changes of the SGNs. For XE991 administration: the 1-month-old *Rest* cKO mice were divided into three groups: the saline, XE991 0.25 mg/kg, and 0.5 mg/kg. XE991 dihydrochloride was obtained from Abcam (Cambridge, UK) and dissolved in 0.9% saline. Mice received a single dose of XE991 (0.25, 0.5 mg/kg) or saline intraperitoneally. ABR was measured in each group of mice before and 30 min after drug administration to characterize the effects of the drug on mouse hearing.

### Statistical analysis

All data are presented as the mean±SEM. Statistical analyses were performed using GraphPad Prism 6.0. The student’s *t*-test was used to compare data between two groups, and ANOVA was used for multiple comparisons. Proportions of cells displaying distinct electrophysiological properties were analyzed using Fisher’s exact test or Pearson’s Chi-square test. *, **, and *** indicated statistically significant results compared to the appropriate controls and indicated *P* <0.05, *P* <0.01, and *P* <0.001, respectively.

## Supplemental Figures

**Figure 1-figure supplement.**
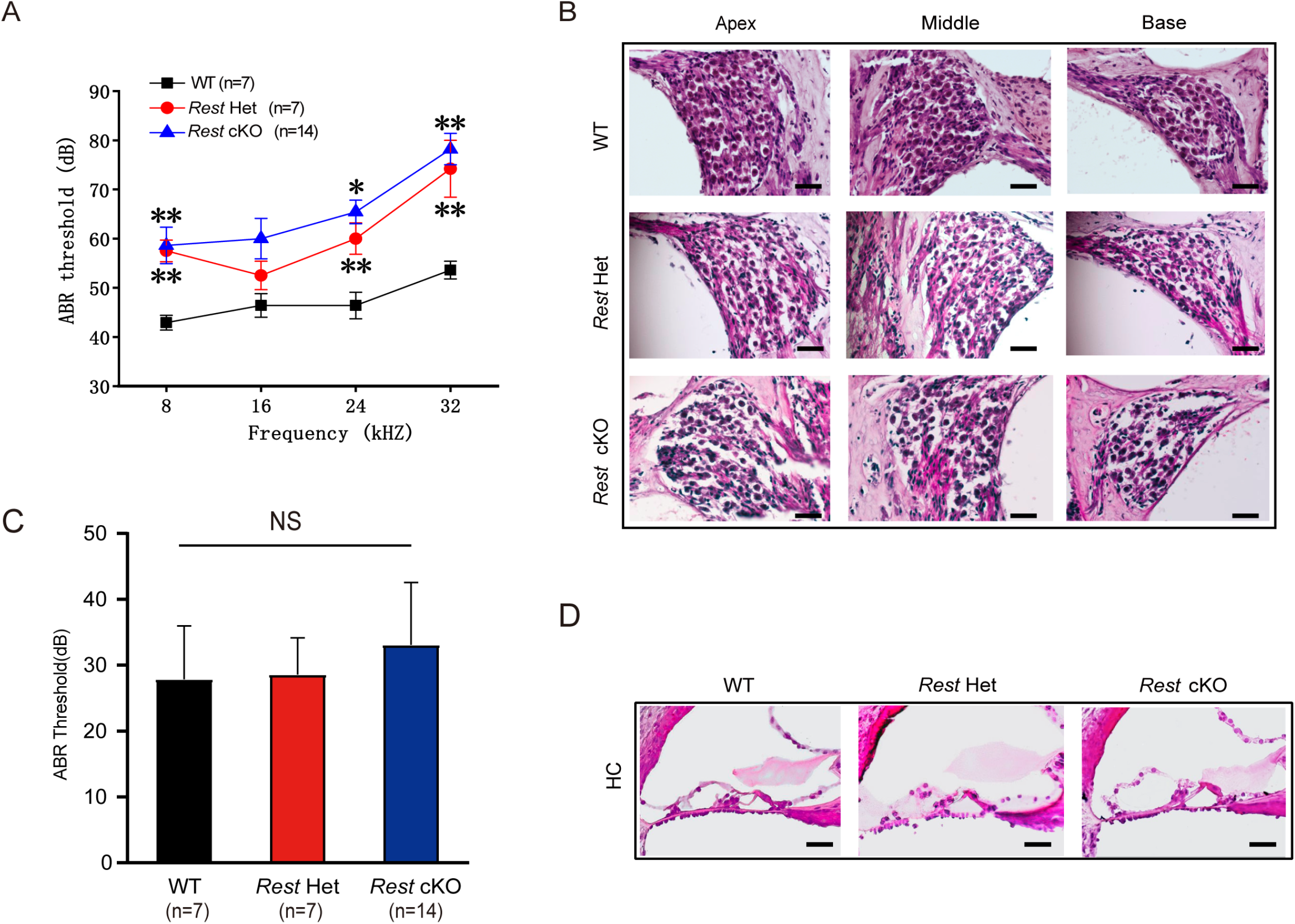
Heterozygotic *Rest* mice show hearing loss. **A, C**, ABR thresholds, and click values were measured in WT, *Rest* heterozygote mice (*Rest* Het), and *Rest* cKO mice at 1-2 months old. **B**, **D**, Morphometry changes in SGNs (B) and HCs (D) were observed in WT, *Rest* Het, and *Rest* cKO mice. Scale bar: 50 μm in B and D. Data are means±SEM, *p < 0.05, **p<0.01.

**Figure 3-figure supplement 1.**
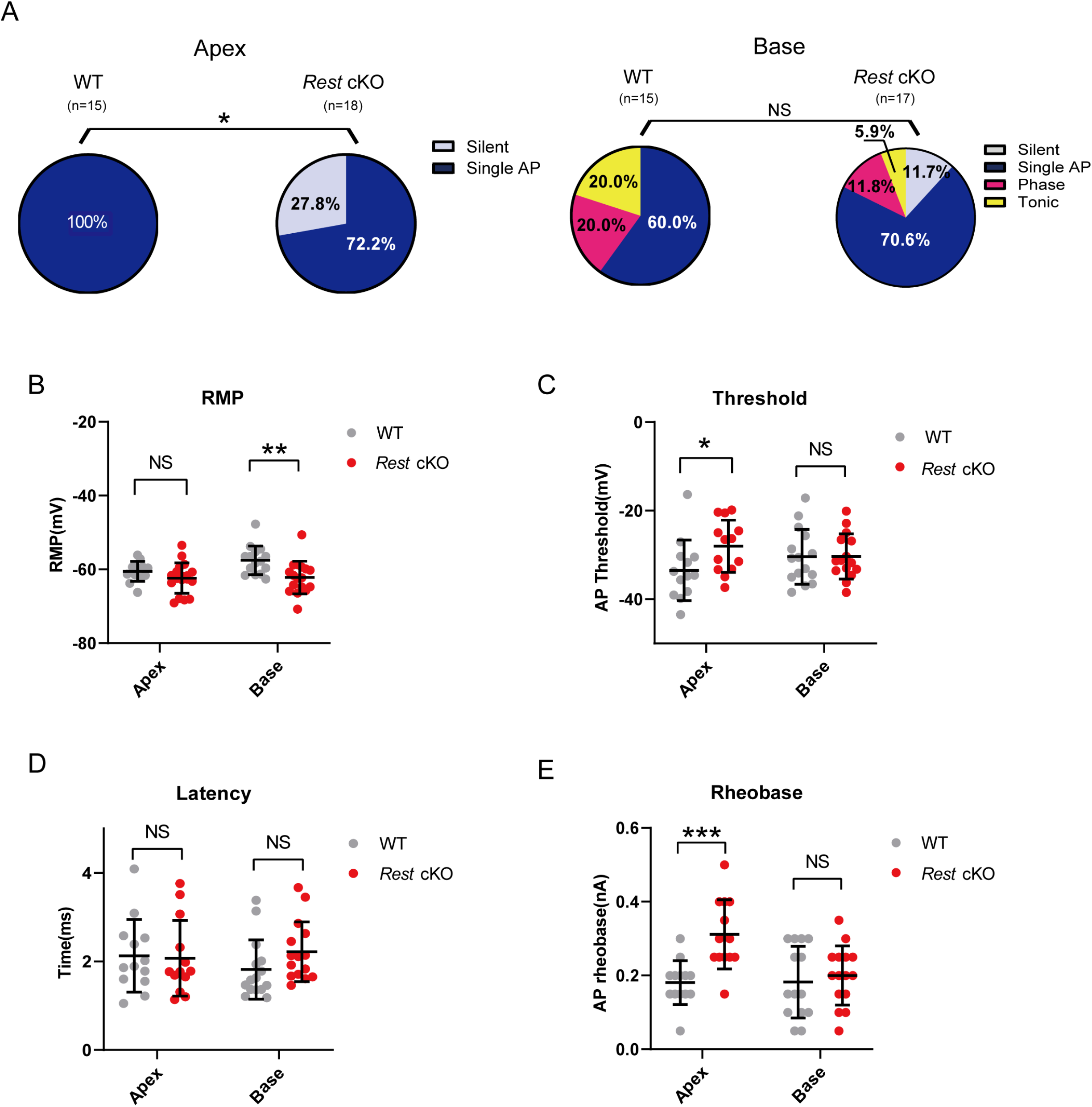
Excitability of SGNs from WT and *Rest* cKO mice at P14. **A,** Pie charts illustrating the percentage abundance of the different spike patterns of SGNs in WT and *Rest* cKO mice at P14. **B–E,** RMPs (B), action potential thresholds (C), latencies (D), and rheobases (E) were recorded from apical and basal SGNs in WT and *Rest* cKO mice at P14. Data are means ± SEM, *p < 0.05, **p<0.01,***p<0.001.

**Figure 5-figure supplement 1.**
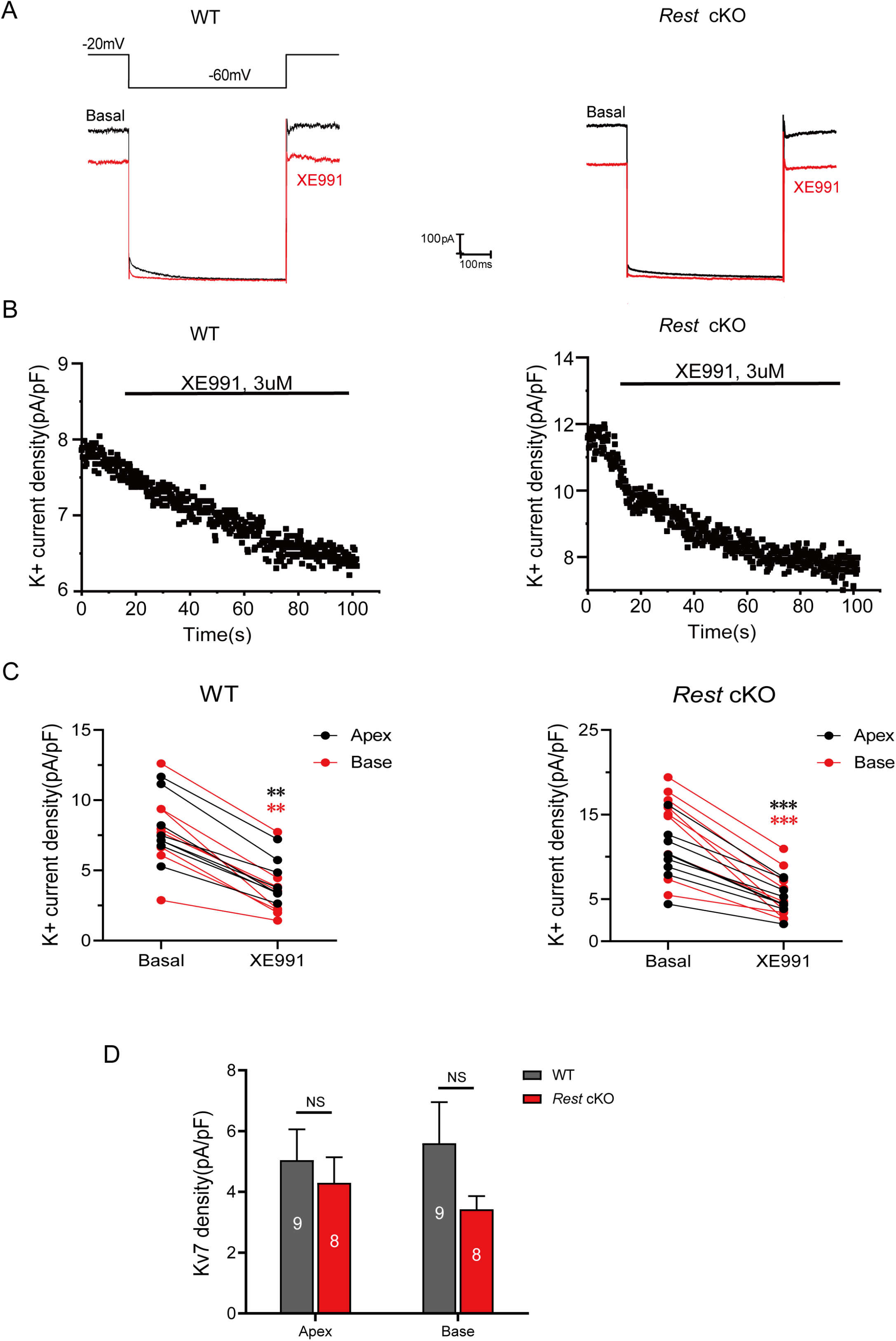
K_v_7 currents recorded from the SGNs of P14 mice. **A**, Representative trace of K_v_7 currents (XE991-sensitive currents) recorded from the SGNs of WT and *Rest* cKO mice at P14. **B**, Time course of the outward K^+^ current inhibition by XE99 1(3μM). **C**, Comparison of K^+^ currents in individual SGNs recorded at −20 mV in the absence (basal) and presence of XE991 (3μM). **D,** Bars show the K_v_7 current density in the apical and basal SGNs from WT and *Rest* cKO mice. Data are means±SEM, **p<0.01, ***p<0.001.

**Figure 5-figure supplement 2.**
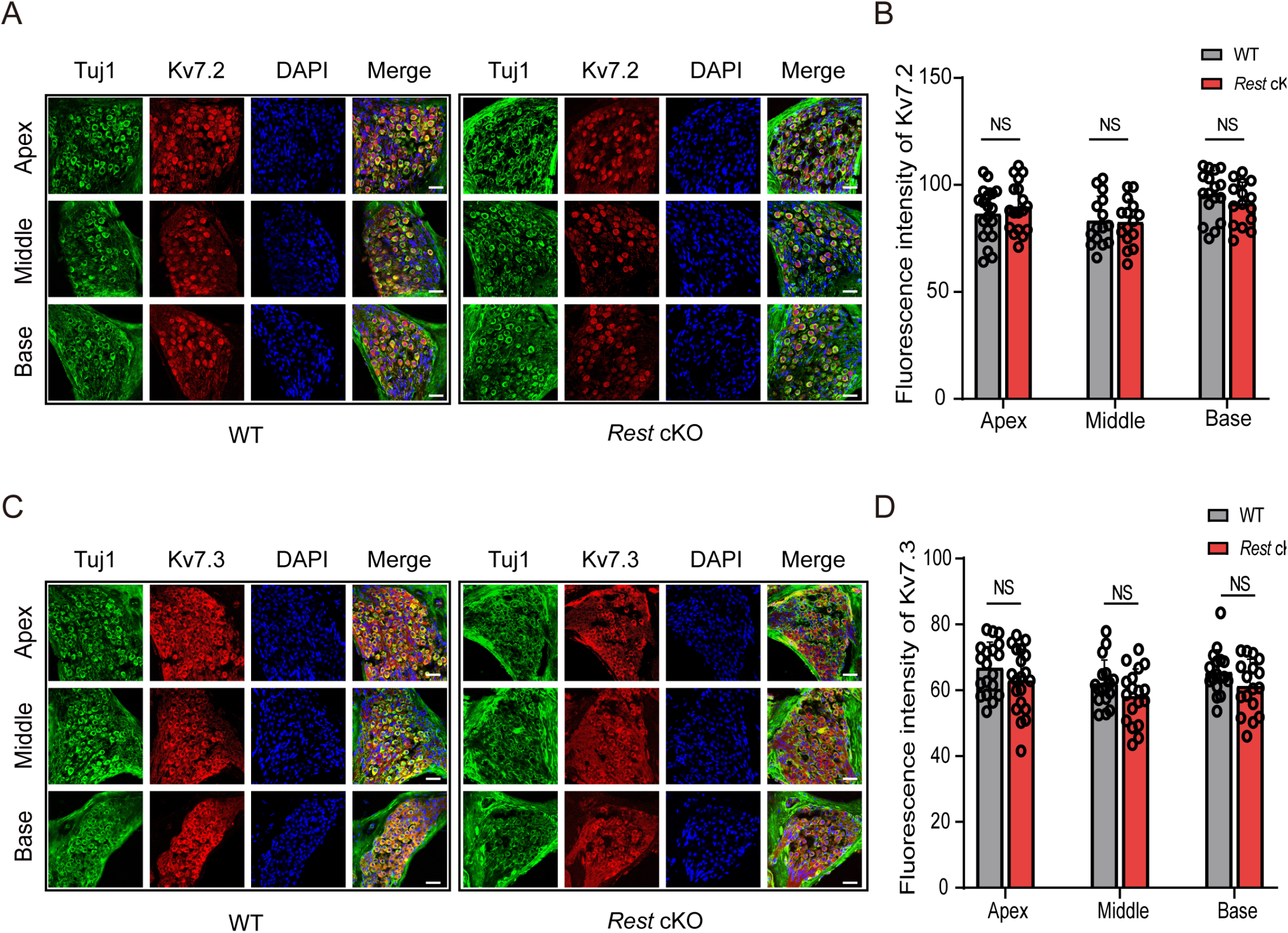
Expression of K_v_7.2 and K_v_7.3 in the SGNs of WT and *Rest* cKO mice. **A**, Expression of K_v_7.2 (red) in the SGNs (green) of 1-month-old WT and *Rest* cKO mice. **B**, Quantification of K_v_7.2 expression in the SGNs of WT and *Rest* cKO mice. **C**, Expression of K_v_7.3 (red) in the SGNs of 1-month-old WT and *Rest* cKO mice. Scale bar: 20μm in A and C. **D**, Quantification of Kv7.3 expression in the SGNs of WT and *Rest* cKO mice.

**Figure 5-figure supplement 3.**
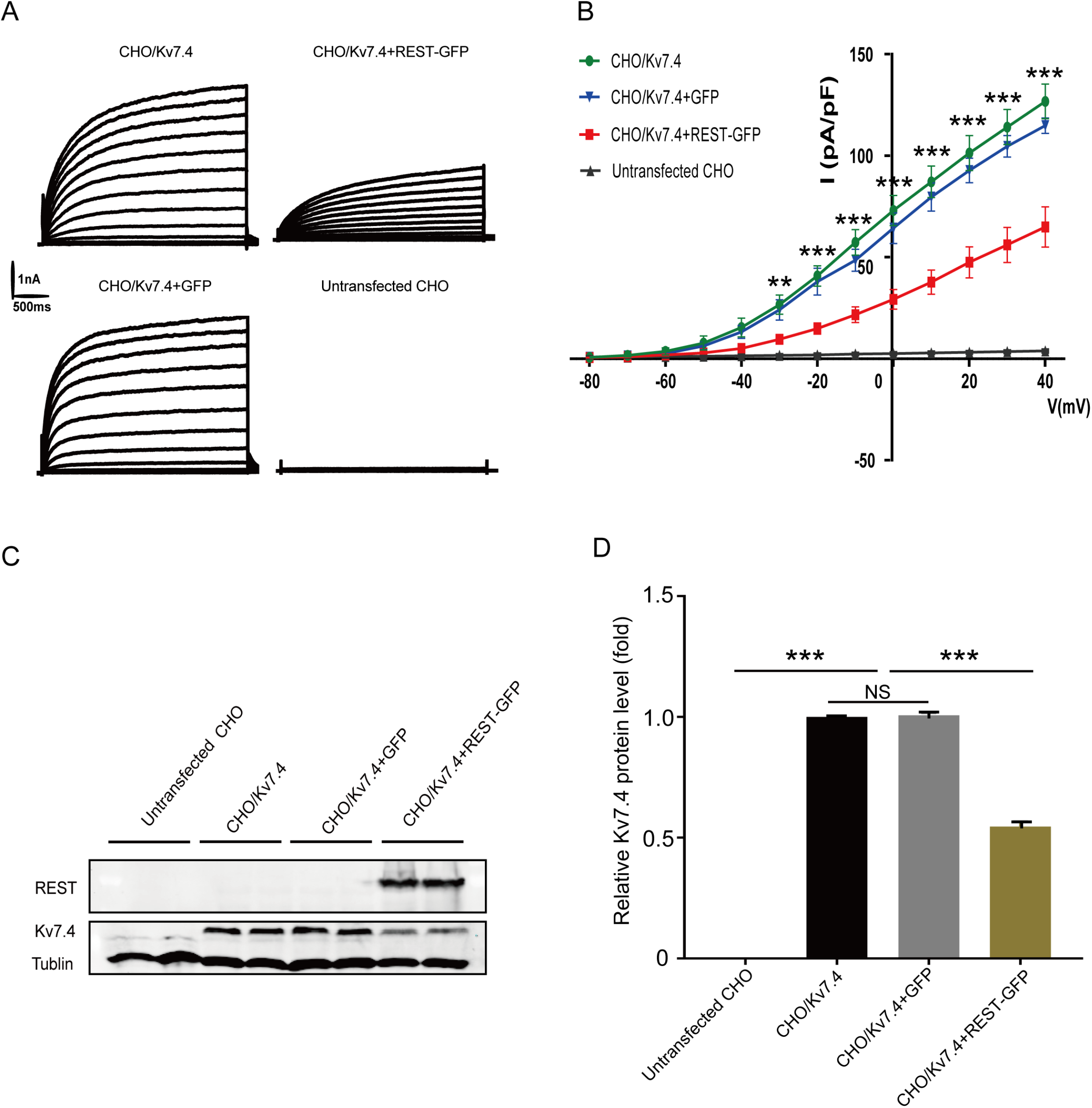
REST inhibited K_v_7.4 channels in K_v_7.4-transfected CHO cells. **A,** Representative current traces were recorded from CHO cells. Cell types included: untransfected CHO cells, CHO cells stably expressing the K_v_7.4 channel (CHO/K_v_7.4), CHO/K_v_7.4 cells that were transfected with the REST-GFP plasmid (CHO/K_v_7.4+REST-GFP), or transfected with GFP only (CHO/K_v_7.4+GFP). Cells were held at −80 mV and stepped up to +40 mV in 10 mV increments. **B**, Corresponding current density-voltage relationships of the cells described in A. **C**, Western blotting of K_v_7.4 expression in the untransfected CHO, CHO/K_v_7.4, CHO/K_v_7.4+REST-GFP, and CHO/K_v_7.4+GFP groups. **D**, Summary data in C show that K_v_7.4 expression is inhibited by transfection with REST. Data are presented as mean±SEM, ***P<0.001.

**Figure 5-figure supplement 4.**
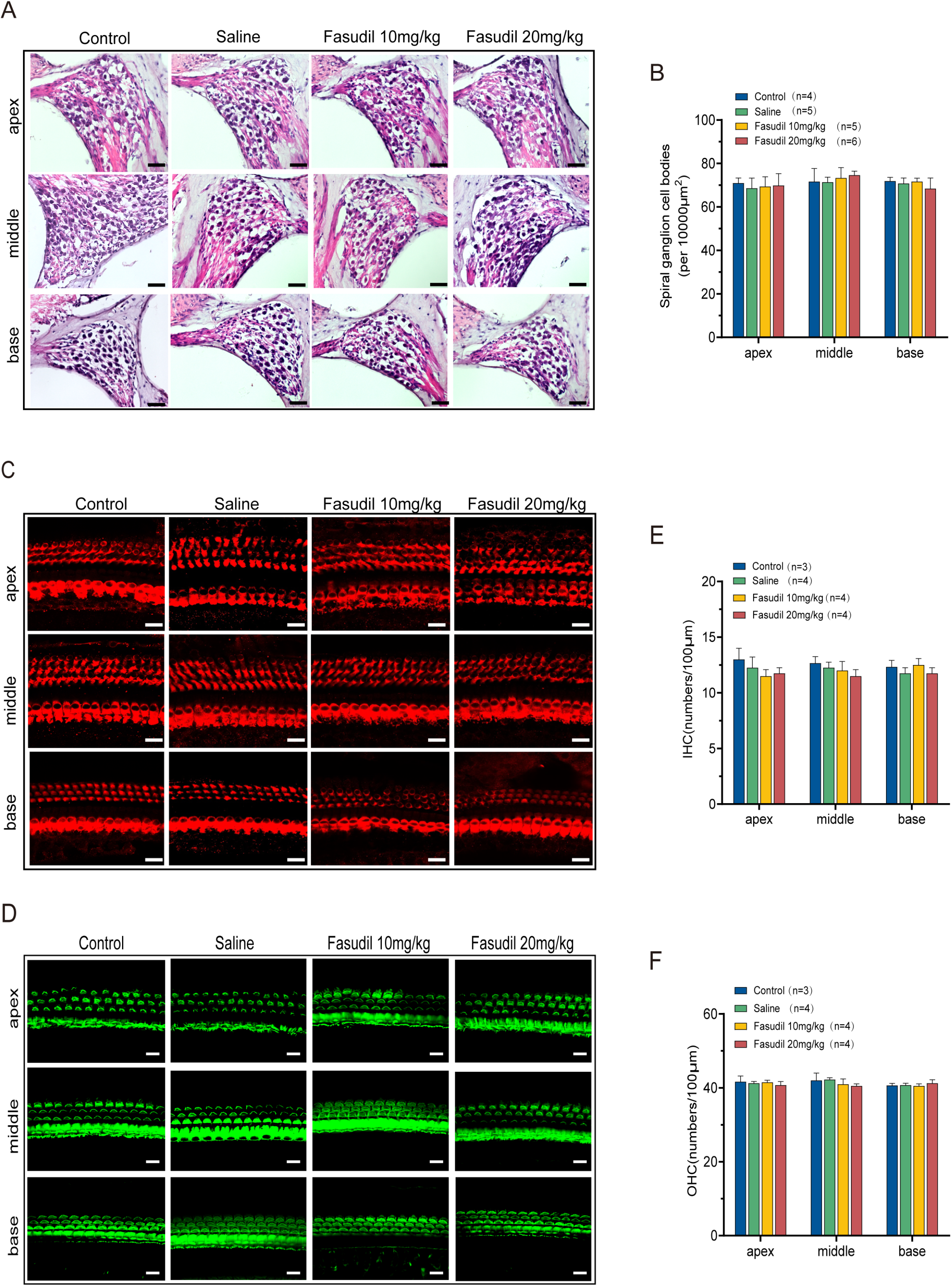
Fasudil did not alter the morphometry of SGNs and HCs. **A,** Morphometry of SGNs in the control, saline, and fasudil (10 mg/kg, 20 mg/kg) treatment groups. Scale bar: 50μm. **B,** Quantification of SGNs in the apical, middle, and basal regions of the cochlea of the different groups. **C, D,** Hair cell bodies (C), and bundles (D) are shown in mice from the control, saline, and fasudil (10 mg/kg, 20 mg/kg) treatment groups. Scale bar: 20μm. **E, F:** The numbers of IHCs and OHCs were counted at the apex, middle, and base of the cochlea from the control, saline, and fasudil (10 mg/kg, 20 mg/kg) treatment groups, respectively. Data are presented as mean±SEM.

**Figure 8-figure supplement 1.**
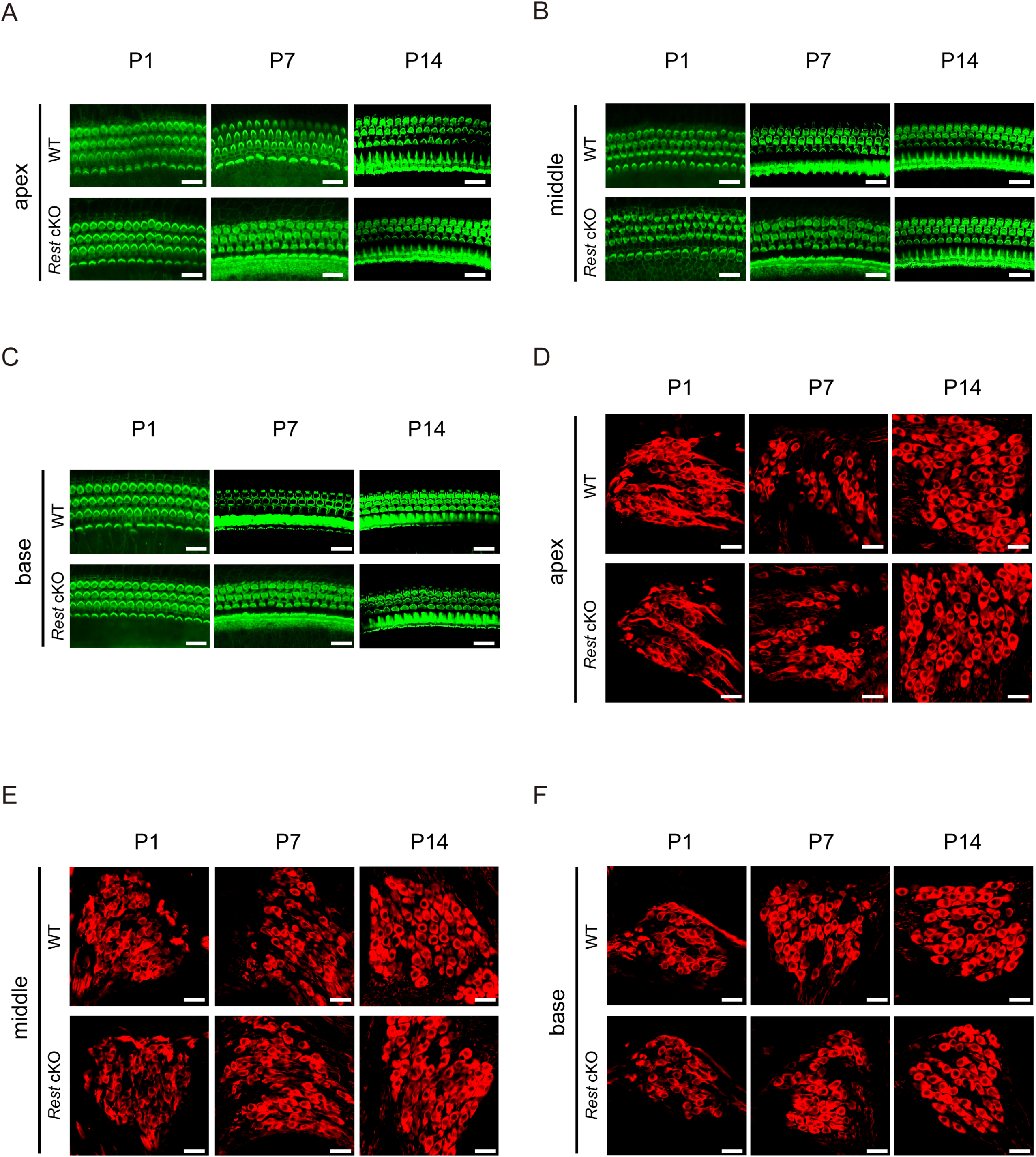
There are no detectable alterations in HC and SGN morphology in *Rest* cKO mice at P1, P7, and P14. **A-C**, Morphology of HCs in the apex, middle, and base of the cochlea of WT and *Rest* cKO mice at P1, P7, and P14. HCs were stained with phalloidin. Scale bar: 20 μm. **D-F**, Morphology of SGNs in the apex, middle, and base of the cochlea of WT and Rest cKO mice at P1, P7, and P14. SGNs were stained with Tuj1. Scale bar: 20 μm.

## Acknowledgments

This work was supported by the National Natural Science Foundation of China (81670939) to PL, by the Natural Science Foundation of Hebei Province of China (H2021206286) to PL, by the Department of human resources and social security of Hebei Province for Talents (A202005003) to PL, by the central government guiding local funding projects for scientific and technological development(216Z7701G) to PL, and by the Science Fund for Creative Research Groups of Natural Science Foundation of Hebei Province ( H2020206474) to PL. ENY was supported by grants from the National Institutes of Health (P01 AG051443, R01 DC015135, R01 DC016099, and R01 AG060504-01). NG was supported by the BBSRC International Partnering Award BB/R02104X/1 and the 100 Foreign Experts of Hebei Province program.

## Author Contributions

PL and ENY designed the research, wrote, reviewed, and edited the manuscript; HZ, HL, ML, SW, XM, FW, JL, XL, HY, FZ, and HS performed the research and analyzed data. NJB and NG reviewed and edited the manuscript.

